# Fluid mechanics of luminal transport in actively contracting endoplasmic reticulum

**DOI:** 10.1101/2023.01.11.523552

**Authors:** Pyae Hein Htet, Edward Avezov, Eric Lauga

## Abstract

The Endoplasmic Reticulum (ER), the largest cellular compartment, harbours the machinery for the biogenesis of secretory proteins and lipids, calcium storage/mobilisation, and detoxification. It is shaped as layered membranous sheets interconnected with a network of tubules extending throughout the cell. Understanding the influence of the ER morphology dynamics on molecular transport may offer clues to rationalising neuro-pathologies caused by ER morphogen mutations. It remains unclear, however, how the ER facilitates its intra-luminal mobility and homogenises its content. It has been recently proposed that intra-luminal transport may be enabled by active contractions of ER tubules. To surmount the barriers to empirical studies of the minuscule spatial and temporal scales relevant to ER nanofluidics, here we exploit the principles of viscous fluid dynamics to generate a theoretical physical model emulating *in silico* the content motion in actively contracting nanoscopic tubular networks. The computational model reveals the luminal particle speeds, and their impact in facilitating active transport, of the active contractile behaviour of the different ER components along various time-space parameters. The results of the model indicate that reproducing transport with velocities similar to those reported experimentally in single particle tracking would require unrealistically high values of tubule contraction site length and rate. Considering further nanofluidic scenarios, we show that width contractions of the ER’s flat domains (perinuclear sheets) generate local flows with only a short-range effect on luminal transport. Only contractions of peripheral sheets can reproduce experimental measurements, provided they are able to contract fast enough.

## I. INTRODUCTION

The mammalian endoplasmic reticulum (ER) is the single largest intracellular structure (see sketch in Fig. 1a). The organelle is made up of membranous sheets interconnected with the nuclear envelope and branching out into a planar network of tubules extending throughout the cell periphery (Fig. 1b) [1]. The ER dynamics on a second scale include the cytoskeleton-assisted tubular network restructuring [2, 3] and an interconversion between two distinct forms, including a narrower form covered by membrane curvature-promoting proteins [4]. The ER morphology and its dynamics presumably enable and facilitate its functions: the ER is responsible for the production, maturation and quality-controlled folding of secretory and membrane proteins, which constitute approximately a third of the cell’s proteome [5]. The organelle’s membranes also harbour the lipid biosynthesis machinery, while its lumen stores calcium. The contiguous nature of the ER is believed to ensure an efficient delivery of all these components across the cell periphery. In particular, ER luminal continuity and transport were demonstrated to kinetically limit calcium delivery for local release [6]. The sensitivity of neurons with long axonal extensions to ER defects in ER morphogens suggests that a perturbed ER transport may link ER integrity and neurodegeneration. Such a link might help explain why mutations in genes involved in ER shaping cause neuronal diseases, including motor neuron degeneration of hereditary spastic paraplegia [7], sensory neuropathy [8] and retinitis pigmentosa [9].

**FIG 1:**
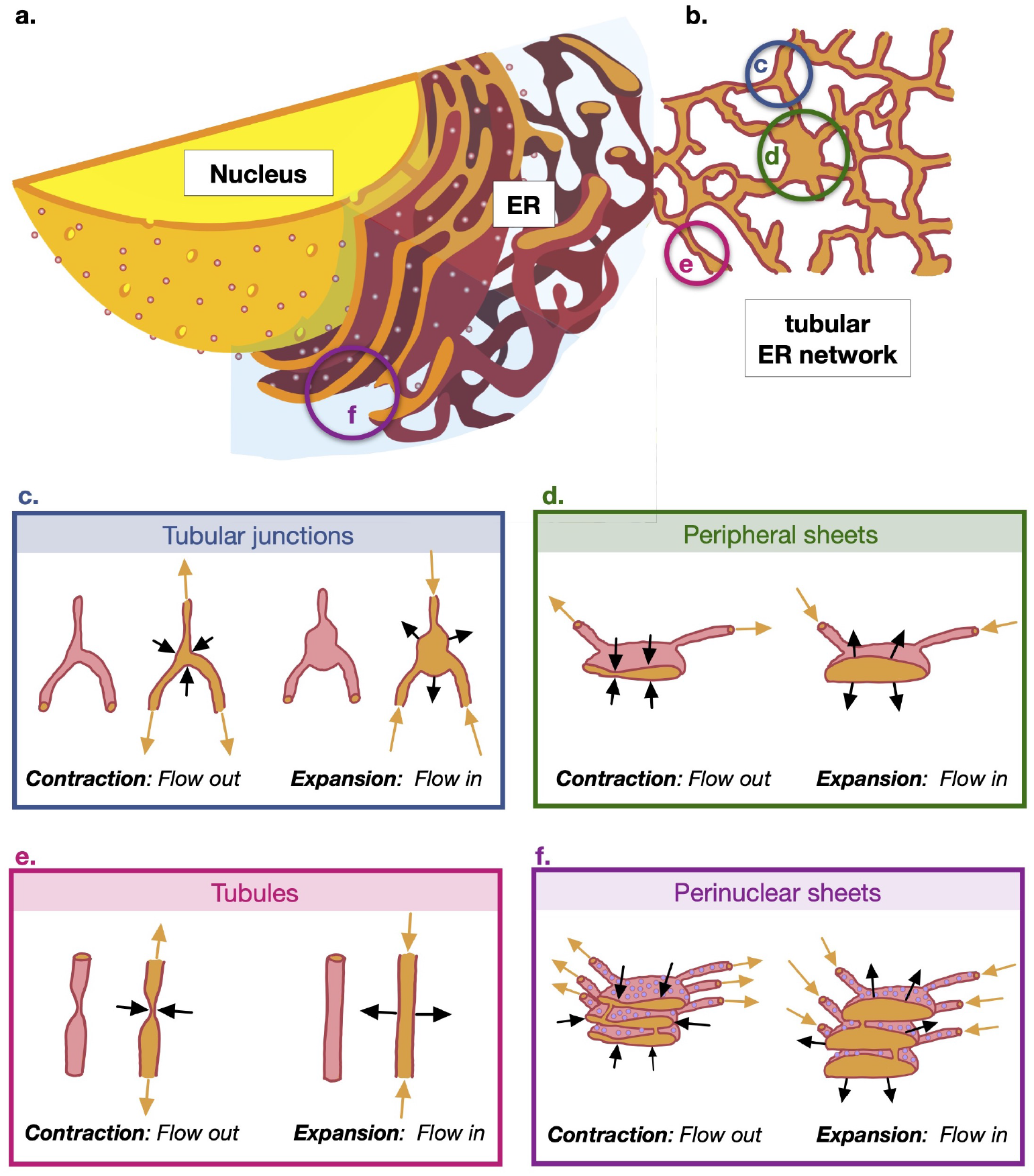
Sketch of the cellular geometry with nomenclature of the sub-cellular structures discussed in the paper. **a**. Cross section of cell showing nucleus and Endoplasmic Reticulum (ER) (adapted from image in public domain [40]). **b**. Cut through cross section of the tubular ER network at the edge of cell. **c**. Sketch of the contraction and expansion of the tubular junctions (3D view and cross section); contractions leads to flow leaving the junction into the network while expansions lead to flow leaving the network and entering the junction. **d**. Contraction and expansion of the peripheral sheets. **e**. Contraction and expansion of the tubules driven by pinching (3D view and cross section). **f**. Contraction and expansion of the perinuclear sheets.

Timely transport of the content within the ER is therefore integral to the function of the cell. The geometry and dimensions of several cell types with extensive ER-containing projections (e.g. neurons and astrocytes) pose a kinetic challenge for material distribution with physiological timing. These considerations predict the need for an active luminal transport to ensure timely material homogenisation across the vast ER. Empirically, the active nature of the ER luminal transport is suggested by a series of observations which include the sensitivity of GFP bulk mobility (measured by FRAP and photoactivation) to ATP depletion [10, 11]. However, these bulk fluorescence intensity dynamics techniques do not provide information on transport mode. By default, the intensity dynamics were historically fitted to diffusion models and the mobility kinetics was often expressed in terms of effective diffusion coefficients. Measurements of single particle motion and chasing locally photoactivated luminal protein marker over distances, circumvent this limit and are also inconsistent with molecular diffusion [12].

However, the mechanism for generating ER luminal flows remains unclear. Understanding the mode of material exchange across the organelle is crucial for rationalising the ER shaping defects-related neuronal pathologies [13], identifying factors controlling ER transport and informing the development of ER transport modulation approaches with health benefits. Based on ER marker velocity fluctuations measured in single particle tracking and the detection of transient narrowing points in the tubules by improved super-resolution and electron microscopy (EM) [11], it has been postulated that these active flows may result from the stochastic contractility (pinching and un-pinching) of ER tubules at specific locations along their lengths (see sketch in Fig. 1e); other plausible mechanisms for flow generation were also considered. However, measurements for testing this pinching hypothesis are currently inaccessible, due to limitations in space-time resolution of live cell microscopy; the live cell-compatible super-resolution techniques achieve resolution of ∼80 nm at the relevant speed, while the tubular radius is estimated in the range 30 − 60 nm [12, 14, 15]. Improvement or resolution currently can only be achieved by trading off speed. To circumvent these experimental difficulties, in the current study we use mathematical modelling to quantitatively analyse the relevant scenarios of actively contractility-driven flows and to explore how various sets of spatio-temporal parameters of ER contractility may produce flows facilitating solute transport in quantitative agreement with experimental measurements.

We illustrate as a simple schematic in Fig. 1 the different contractility mechanisms in the different regions of the ER and where they are located in relation to the cell centre/nucleus. The ER close to the cell nucleus is geometrically complex, consisting of stacks of perinuclear sheets (Fig. 1a). Away from the nucleus, the ER geometry simplifies considerably and the ER at the cell periphery is comprised of a planar network of tubules (Fig. 1b). In the current work we study flows and transport in this planar, tubular region; this is also the region of the ER network in which single particle tracking measurements were carried out in Ref. [11]. We consider as potential driving mechanisms for the observed solute transport the contractility of components located in the same planar tubular region, namely, tubular junctions (Fig. 1c), peripheral sheets (Fig. 1d), and tubules (Fig. 1e), as well the contractility of perinuclear sheets (Fig. 1f) located closer to the nucleus.

First, assuming that the flow is driven by tubule contractions (shown schematically in Fig. 1e), we construct a physical and mathematical model of the ER network and solve it for the flows inside the network (description in Methods §IV). We use our model to carry out numerical simulations to study the motion of Brownian particles carried by these flows and show that the tubule pinching hypothesis is not supported by the results of our model (§II A), a result independent of the network geometry (§II B). The failure of active pinching to drive strong flows can be rationalised theoretically by deriving a rigorous upper-bound on the rate of transport induced by a single pinch, (§II C), including possible coordination mechanisms (§II D). Only by increasing both the length of pinches and their rate can beyond admissible values can we produce particle speeds in agreement with measurements (§II E). We then explore in §II F two different hypotheses as possible explanations for the active ER flows, first the pinching of the junctions between tubules (Fig. 1c) and then the pinching of the two types of ER sheets, perinuclear (Fig. 1f) and peripheral (Fig. 1d). We investigate the conditions under which these contractility mechanisms would be consistent with the experimental measurements of active transport in Ref. [11].

The question of how the endoplasmic reticulum homogenises its content across cell expanses is an open problem which extends to the fundamentals of cell biology. Even though it is highly debated in the field, it is extremely difficult to study due to the enormous technical limits of directly observing intraorganellar nanofluidics. Thus, our study represents a meaningful effort to break through this impasse by conducting a meticulous analysis of nanofluidic scenarios. The findings we present constitute the best currently possible endeavour to shed light on this challenging problem. Our results do suggest that the biological origin of solute transport in ER networks remains open and call for extensive empirical exploration of the alternative mechanisms for flow generation.

## II. RESULTS

### A. Tubule contractility-driven ER luminal motion yields inadequate transport kinetics

To assess the kinetics of particle motion in the lumen of tubular structures, detailed in Fig. 1, in response to their contractility, we generate an *in silico* simulation model of the process. The model incorporates local calculations for the low-Reynolds number hydrodynamics of a contracting tubule, assuming in the first instance the no-slip boundary conditions at the tubule walls (i.e. Poiseuille hydrodynamics), into a global analysis of the flows throughout the network geometry, by using Kirchhoff’s Laws and standard graph theoretical results (see Methods §IV for details).

We initially implement the model’s numerical simulations of particle transport in a reconstructed ER network of a COS-7 cell [11] (which we label C0, see §IV A) with tubules locally contracting stochastically according to the spatiotemporal parameters suggested by microscopy measurements [11]. We will therein refer to these contractions, illustrated in Fig. 1e, as ‘pinching’, with relevant parameters that include the duration and frequency of pinch events, and the length that the pinch sites occupy along the tubule (for details of the pinching kinematics, see Methods §IV B). An estimate reveals that these pinches are indeed afforded by biologically realistic forces, of the order of 30 pN (see Methods §IV H).

The fluid flows in the edges of the network (model in §IV C computed as detailed in §IV D) reveal a rapid direction alternation of luminal currents (on average with a frequency of approximately 50 Hz), as reflected in the changes of the axial velocity sign (Fig. 2a), with an average flow speed of 1.3 μm/s (see also Supplementary Video S1). Further, the resulting instantaneous speed distribution of Brownian particles advected by these flows (methodology in §IV E) is considerably shifted relative to the experimental counterpart [11] towards lower values (Fig. 2b). Similarly, the distribution of average edge traversal speeds (defined precisely in §IV F) from our simulations (mean 4.4 μm/s, solid blue line in Fig. 2c) is lower than the experimentally measured speed distribution (mean 45 μm/s; Fig. 2c, inset) by an order of magnitude.

**FIG 2:**
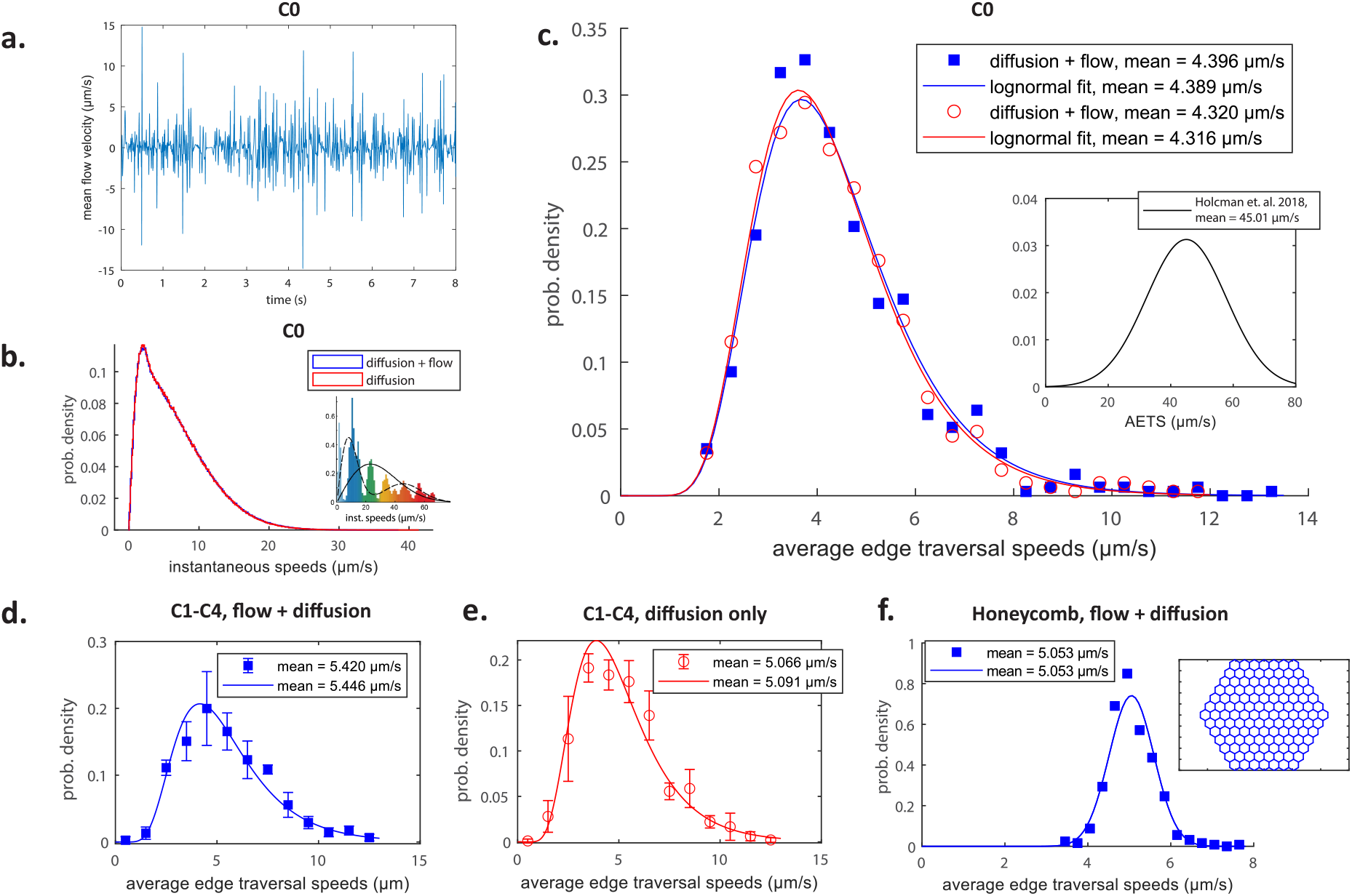
A quantitative test of the pinching-tubule hypothesis. **a**. Cross-sectionally averaged flow velocities in a typical edge as obtained in our simulations. **b. c**. Histograms of instantaneous speeds (b) and edge traversal speeds (c) using data from simulations in the C0 network with flow (blue) and with just diffusion (red). The insets in (b) and (c) illustrate the distributions of instantaneous speeds and average edge traversal speeds respectively as experimentally measured in Ref. [11]. The symbols indicate the values taken by the probability mass function and the curves are log-normal distributions fitted to all average edge traversal speeds obtained. **d-f**. Histograms of average edge traversal speeds obtained from simulations in networks C1-C4 from Fig. 9d with flow (d) and only diffusion (e) and from simulations in the regular honeycomb network with active flows (f). The inset in (f) illustrates the honeycomb geometry. Points indicate mean ± one standard deviation over the four networks (C1-C4) of normalised frequencies in each speed range; curves are log-normal (d-f) or normal (f) distributions fitted to all average edge traversal speeds for each set of pinch parameters. The means of the original simulation results and of the fitted distributions are indicated in the legends in each of (c-f).

Moreover, the results for all measures of transport under pure diffusion, in the absence of pinching-induced flows, are virtually indistinguishable from those where the pinching-driven flows are included (see Fig. 2b-c). Within the framework of transport theory, this conclusion can be rationalised by estimating the relevant Péclet numbers that measure the relative importance of advection and diffusion. Using the average value 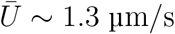 of the mean flow speeds over time and edges as a velocity scale, we may estimate a mean Péclet number as Pe ∼ 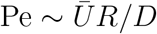. Using *R* = 30 nm and the measured diffusivity *D* ≈ 0.6 μm^2^s^−1^ [11], this leads to the estimate Pe ∼ 0.07. Flows affect transport for Péclet numbers above order-one values [16], and therefore the pinching-driven flows have a negligible influence on the transport inside the ER network. In order for fluid motion to have a noticeable effect, one would thus need flows to be either stronger, or sustained in one direction for longer.

The chaotic flows produced by the pinching events with stochastic parameters suggested by resolution-limited microscopy [11] appear too weak to generate fast, non-diffusive edge traversals.

Importantly, our conclusions remain unchanged upon relaxing the no-slip boundary conditions, an important point to consider since the tubule membranes themselves could be moving in response to the nanoscale fluid flow. This can be modelled by introducing a finite slip boundary condition to couple the membrane-bound lipids with luminal flows (see Methods §IV G for details). Our simulations show that slip boundary conditions have virtually no effect on the average edge traversal speeds (Fig. 3a). Physically, while a non-zero wall slip does change the shape of the flow profiles in the tubules (Fig. 3b), it does not modify the mean flow speeds which directly affect global particle transport. These results justify our use of Poiseuille hydrodynamics (no-slip boundary conditions) throughout this work.

**FIG 3:**
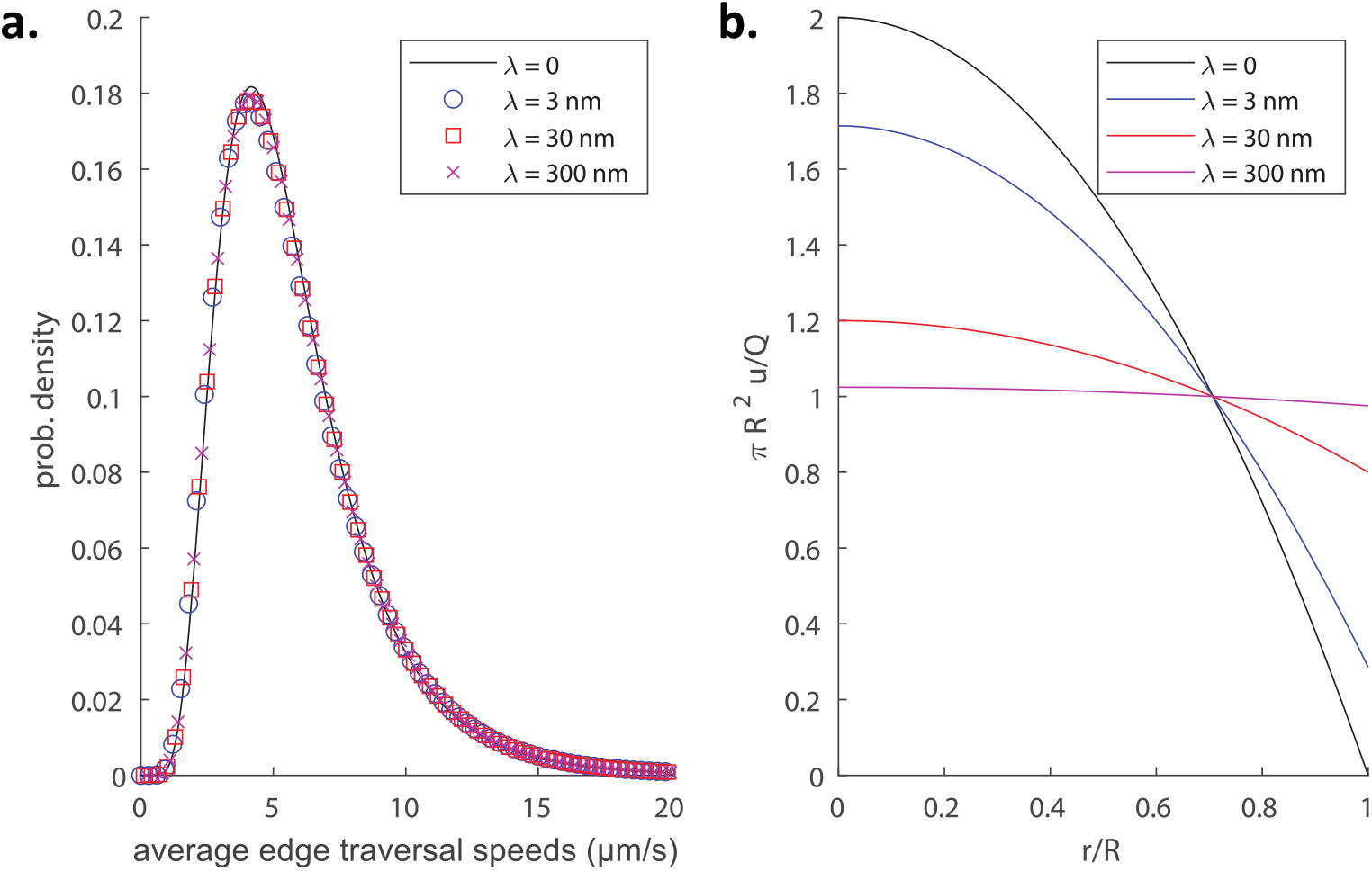
Impact of non-zero slip length on transport and flow. (a) Distributions of average edge traversals speeds in simulations of a C1 network pinching with the original pinch parameters as measured in Ref. [11] and used in Fig. 2, for different slip lengths *λ* = 0, 3, 30 and 300 nm. (b) Longitudinal flow profile *u*(*r*) inside a cylindrical tubule for different slip lengths, all with the same volume flux *Q*; an increase of the slip length leads to a redistribution of the flow in the cross section.

### B. Tubule pinching-induced transport is network geometry independent and fails to facilitate luminal homogenisation

To estimate the contribution of the network geometry to the outcomes of our transport simulations, we compare results across four different ER structures, which we label C1-C4. We illustrate these reconstructed networks along with the source data in Fig. 9. The distributions of the average edge traversal speeds appear insensitive to ER structure variations for both pinching-induced and exclusively diffusional transport. This is reflected in the small deviations from the mean of the data points averaged across the different structures (Fig. 2d-e). Further, pinching-induced flows inside a regular honeycomb network (Fig. 2f, inset) with a typical ER edge length (1 μm) appear to be within a reasonable variance compared to the natural networks. Therefore, this mathematical idealisation of the ER network geometry can be used for exploring the consequences of the network ultrastructure contractility on transport kinetics in a standardised manner.

To test the effectiveness of the particle velocities for facilitating luminal material exchange across the ER, we track homogenisation kinetics by measuring intermixing of particles of two distinct colours equally seeded in each half of a honeycomb network at *t* = 0 (Fig. 4; see also Supplementary Video S2 for the flows inside such a network driven by pinches with the parameters from [11]). The measure of homogenisation is given by the variance Var(*ϕ*(*t*)) ≡ Var(*n*_*b*_(*t*) − *n*_*r*_(*t*)) over twenty regions of the network (Fig. 4a, horizontal lines) of the difference between the numbers *n*_*b*_ (blue) and *n*_*r*_ (red) of particles of each colour in region; note Var = 0 represents perfect homogeneity. The measures of mixing over time for pure diffusion and active transport with parameters [11] from Fig. 2 show a nearly complete overlap (Fig. 4e; see also Supplementary Videos S3 and S4), distinct from faster mixing under stronger flows in a network driven by pinches whose lengths are increased to their maximum possible value (i.e. the length of the tubule) and which are in addition sped up by a factor of 10 (see Supplementary Video S5; see also §II E for a discussion of the effects of artificially strong pinch parameters on average edge traversal speeds). This indicates that even if the experimental particle speeds were overestimated, the presumed pinching parameters would be inadequate to facilitate ER luminal material exchange.

**FIG 4:**
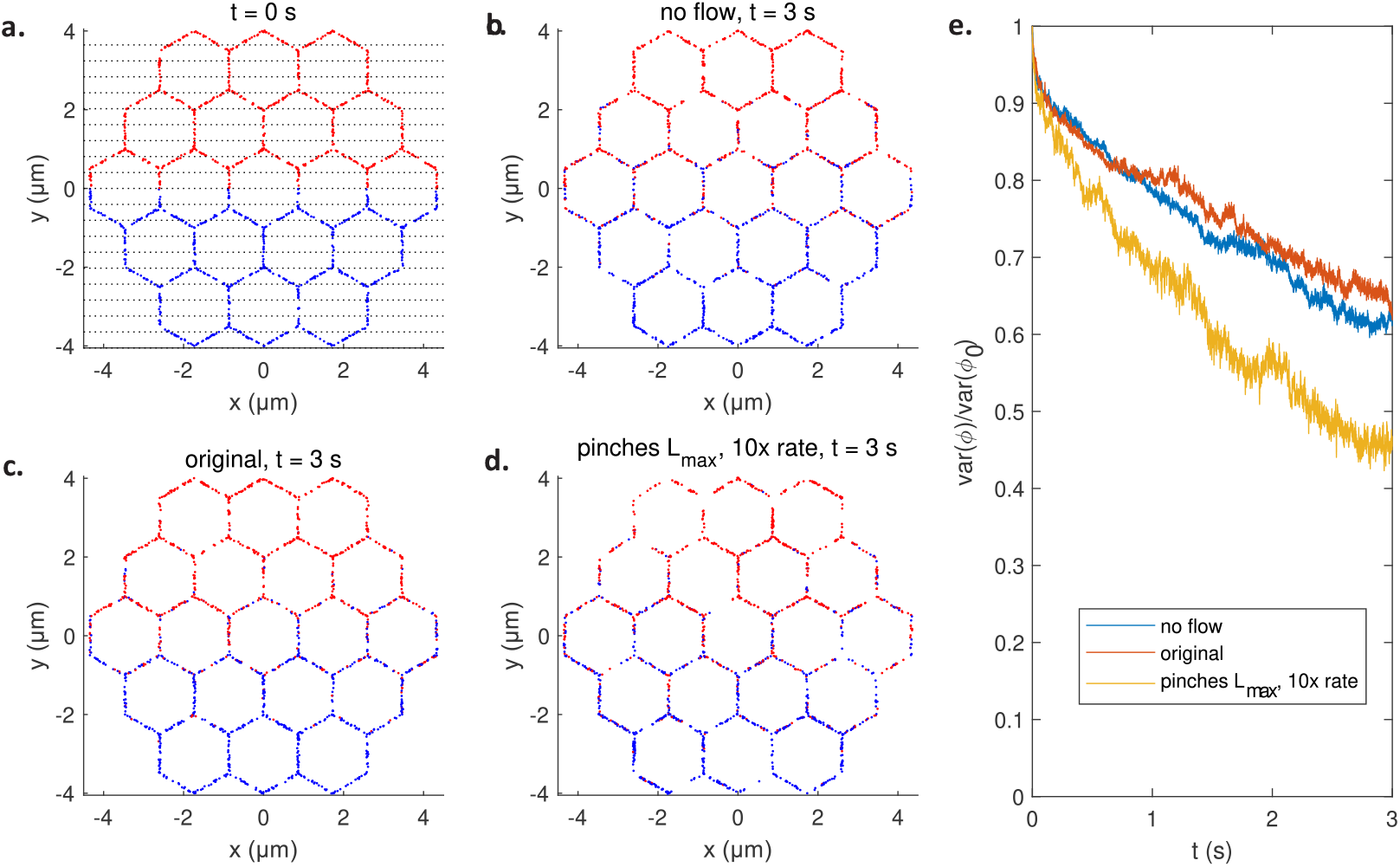
Mixing by active pinching flows. **a**. Initial configuration of blue and red particles in honeycomb network. The strips used to quantify mixing are illustrated in black dotted line. **b-d**. The configuration after *t* = 3 s of mixing in a passive network with no flow (b), an active network pinching with the original pinch parameters (c), and an active network pinching with maximally long pinches at 10 times the original rates (d). **e**. The measure of mixing Var(*ϕ*(*t*)) against *t* for the passive network (blue), the network pinching with the original parameters (red), and the network pinching with maximally long and 10x faster pinches (yellow).

### C. Theoretical analysis of advection due to a single pinch explains weak pinching-induced transport

The slow luminal transport driven by the pinching-induced flows is intrinsically linked to the volume of fluid expelled by each pinch during a contraction. The fundamental reason underlying the weak pinching-induced transport is that individual pinches are very weak generators of flow; even in the best possible configuration, the volumes of fluid pushed by each pinch are too small to impact luminal transport. Specifically, in Methods (§IV I 1), we mathematically show that, given a pinch of length 2*L*, the maximum distance Δ*z*_max_ a suspended particle may be advected instantaneously by an individual pinch is

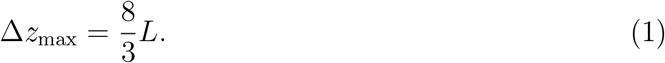

Using the experimentally-measured average value of the pinch length 2*L* = 0.14 μm [11], a typical pinch can then propel a particle by a maximum distance of Δ*z*_max_ ≈ 0.19 μm.

This may, equivalently, be framed in terms of velocities. A transported particle experiences an average velocity during the contraction of at most *V*_max_ = 8*L/*3*T*, where *T* is the duration of a contraction or a relaxation. Using the pinch length as above and the average values of 2*T* = 0.213 s [11] this leads to the estimate *V*_max_ ≈ 3.9 μm/s, which is an order of magnitude smaller than the measured typical edge traversal speed of ∼ 45 μm/s, consistent with the order of magnitude difference between the measurements and the predictions of the computational model.

Note that this theoretical argument relies solely on the magnitude of fluid volumes expelled out of (or driven into) the pinching regions. These volumes are unaffected by the boundary conditions at the tubule walls because the presence of a non-zero slip length only changes the shape of the flow profile and not the volume fluxes (see Fig. 3); any potential slip is thus inconsequential for averaged measures of particle transport.

### D. Marginal addition of coordinated contractility to luminal transport

The theoretical upper bound in the previous section for the maximum luminal transport producible by an individual pinch is realisable only in the hypothetical situation where the flow generated by the pinch is all directed to one end of the tubule i.e. when the other end is effectively blocked. However, content transport produced during the contraction of a single tubule would then be reversed when the tubule relaxes back to its original state, with a single pinch site expected to exhibit only reciprocal (i.e. time-reversible) motions. Any advection contributing to edge traversals must thus be dominated by non-reciprocal motions of multiple pinches resulting in net displacements of solute particles.

The simplest system capable of producing non-reciprocal motions consists of two pinches, and the optimal sequence of motions to maximise the resulting advective particle displacement is illustrated in Fig. 5. We show in Methods (§IV I 2) that this is indeed the optimal two-pinch coordination, which results in a time-averaged displacement equal to the upper bound derived in Eq. (1). Since this optimal sequence of motions involves one pinch site starting a pinch halfway through the pinching of the other site, it is reasonable to estimate its duration as 3*T*, and therefore an average particle speed of 8*L/*9*T* ≈ 1.3 μm/s. The low particle speed achievable by the optimal coordination between two pinches suggests that a very high level of coordination among multiple tens of pinches per tubule would be required to reproduce the measured edge traversals.

**FIG 5:**
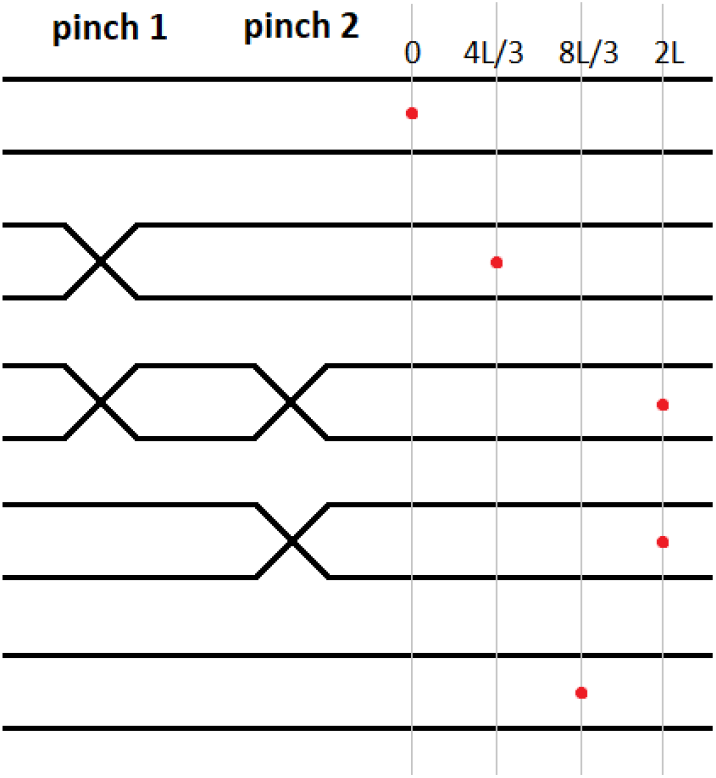
Illustration of coordination mechanism allowing the interactions between two pinch in series to induce the net transport of a suspended particle; the mechanism is akin to small-scale peristalsis.

### E. A combination of high frequency and pinch length is required to replicate experimental particle speeds

Since the magnitude of ER contractility suggested by microscopy [11] (pinch lengths and frequency) does not explain the measured speeds, we set out to explore different sets of parameters that may generate particle velocities closer to the experimental measurements. The currently achievable imaging spatiotemporal resolution limits the detection of tubular diameter contractility by microscopy. Therefore, it is reasonable to postulate that the relevant parameters may have been underestimated. We simulate ER transport varying individually or in combination the values of pinch duration 2*T*, waiting time *T*_*wait*_ between successive pinches on a tubule, and pinch length 2*L*.

We first decrease both the original [11] values of *T* and *T*_*wait*_ by the same factor of 1*/α*, where *α* ≥ 1, and simulated particle transport in the honeycomb network (Fig. 6a). In effect, this simply ‘fast-forwards’ the flows in the original active network by a factor of *α*, and Brownian particles of the original diffusivity are released into this sped-up flow. Instantaneous and edge traversal speeds exhibited corresponding increases when we increased the value of *α* (Fig. 6a). An extreme value of *α* = 100 produces an average edge traversal speed distribution that peaks at around 8 μm/s (Fig. 6a). Similar results are observed in the C0 network from a COS-7 cell (Fig. 6b). The longer tails of these distributions (compared to those from the honeycomb network) result from the variation in edge lengths in the real network, with shorter edges, across which edge traversals are correspondingly fast, contributing to the tails.

**FIG 6:**
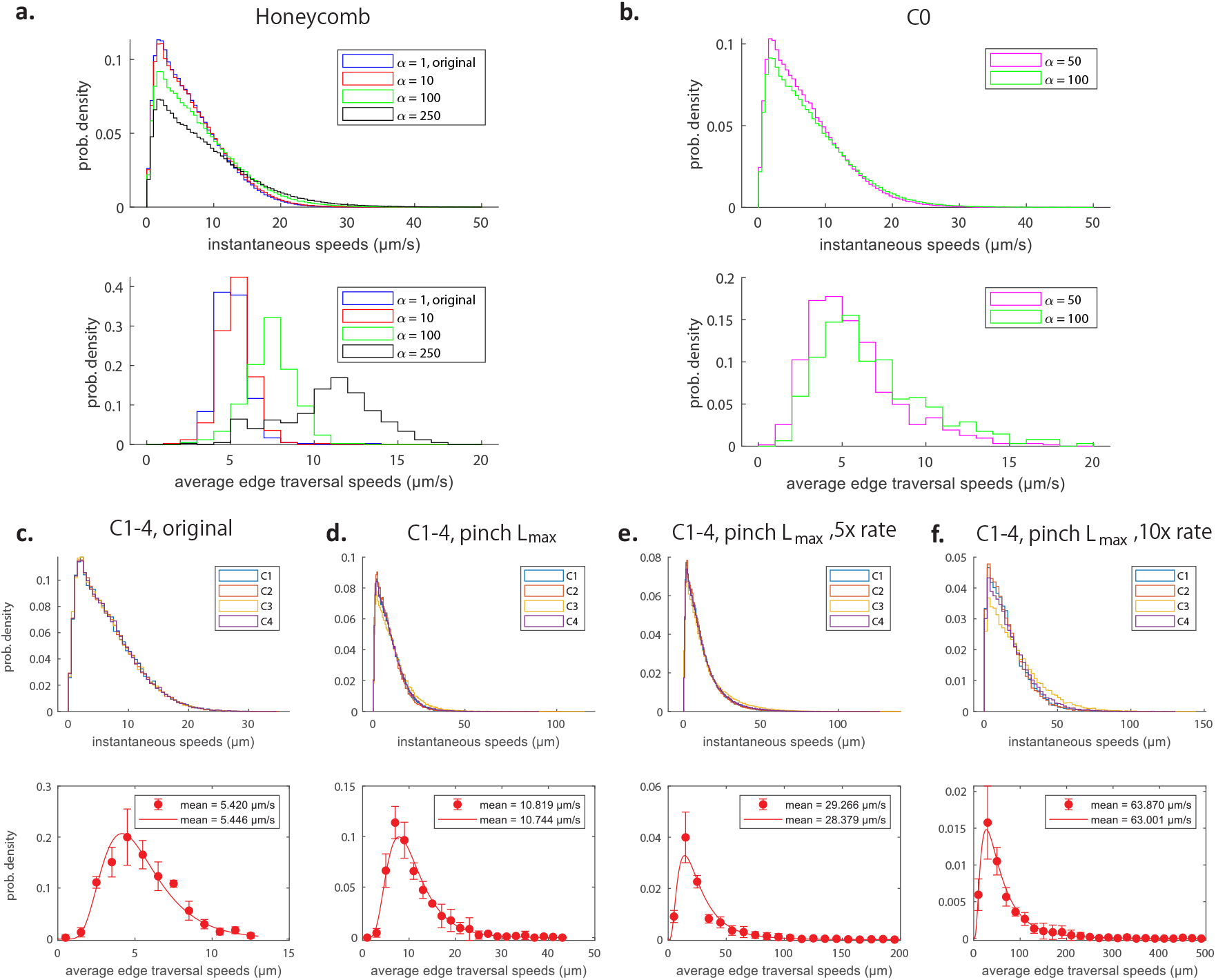
Histograms of instantaneous speeds (top) and average edge traversal speeds (bottom), for **a-b**. an active honeycomb network (a) and the reconstructed C0 network from Fig. 9a-c (b) with pinch parameters 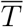 and 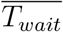 decreased to 1*/α* times the original values from Ref. [11], and the same measured diffusivity *D* = 0.6 μm^2^s^−1^, and for **c-f**. the C1-4 networks from Fig. 9d with varying pinch parameters: original parameters from Ref. [11] (c); pinch length increased to the total length of the tubules (d); a five-fold increase in the rate of pinching and pinch length set to the total length of the tubules (e); a ten-fold increase in the rate of pinching and pinch length set to the total length of the tubules (e). Bottom rows: points indicate mean ± one standard deviation over the four networks (C1-C4) of normalised frequencies in each speed range; curves are log-normal distributions fitted to all average edge traversal speeds for each set of pinch parameters; insets show means of original simulation results and of fitted distributions.

These results suggest that, in order to produce average edge traversal speeds of the same order as the experimental values, we would need an active network sped up by an unrealistic factor considerably greater than 100, probably on the order of *α* ∼ O(10^3^) or higher and corresponding to multiple thousands of pinches occurring on average per second on each tubule. Similarly, it took an extreme increase in pinch site size spanning the entire length of an average tubule, only to yield an average speed of ∼ 10 μm/s (Fig. 6d).

Next, we attempt to obtain a better fit to experimental data by combining changes in both the pinches’ time and geometry parameters. In Fig. 6e-f these maximally long pinches are sped up by a factor of *α* = 5 and 10 respectively, which yields speeds averaging around 30 μm/s and above 60 μm/s respectively. Notably, the tail of the speeds distribution appear longer than that seen in the experiments.

### F. Luminal transport kinetics derived from contractile ER tubular junctions and sheets

As shown above, establishing effective transport in a tubular constrictions-driven model based on realistic ER fluid dynamics requires a set of questionable assumptions, compelling us to explore alternatives. Thus, we set out to investigate how ER luminal transport would be impacted by the contractility of its structural components with volumes larger than tubules: (i) the tubular junctions (Fig. 1c), (ii) the perinuclear ER sheets (Fig. 1f) and (iii) the peripheral sheets (Fig. 1d).

First, we run numerical simulations of transport driven by contracting junctions on an ER network (C1 from Fig. 9d; sketch of junctions in Fig. 1c). Since junctions contracting at the tubular pinch temporal parameters measured experimentally yielded inadequate transport, we consider contractions/relaxations with duration 2*T* exponentially distributed with a mean of *α*^−1^ times the original value in Ref. [11] and the waiting time *T*_*wait*_ between subsequent contractions/relaxations exponentially distributed with a mean *β*^−1^ times the original value (i.e. values of *α >* 1 and *β >* 1 reflecting faster and more frequent pinches). Naturally, the results depend on the choice of the volume Δ*V* expelled by a junction during each contraction. Using fluorescence microscopy images, we may estimate the volume in junctions (see Methods, § IV J 1). We assume that the junction volumes are drawn from a normal distribution with the same mean and standard deviation as our data set i.e. the distribution N(0.0045 μm^3^, 0.0021 μm^3^). Volumes greater than the maximum value in our experimental estimate (0.0081 μm^3^) or less than the minimum value (0.0020 μm^3^) are rejected.

We show the results in Fig. 7a. The thick solid line illustrates a set of values of (*α, β*) which produces distributions of average edge traversal speeds with means similar to the experimental values in Ref. [11], and the dashed lines one standard deviation away. We are interested in values of (*α, β*) close to unity because this corresponds to junctions which pinch with similar pinch durations and frequencies as the experimentally observed tubule pinches and are therefore biologically plausible; (*α, β*) ≈ (2.5,2) is the closest pair on this line to unity. However, only for *α* ≈ 1 we obtained average edge traversal speed distributions of reasonable shapes (i.e. approximately Gaussian), which would require a large *β* = 5 to match experimental results quantitatively.

**FIG 7:**
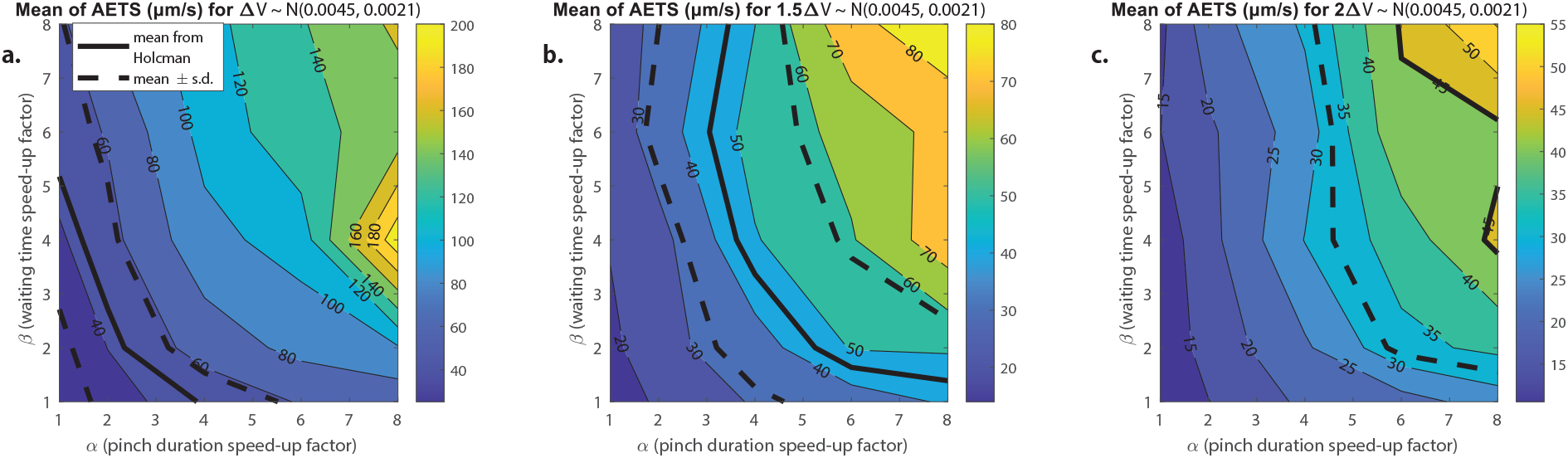
Contour plots of the mean values of the average edge traversal speeds obtained from simulations of our model in a junction- and tubules-driven C1 network from Fig. 9 with different values of (*α, β*) and with contraction volumes Δ*V* expelled during each contraction drawn from: **a**. the normal distribution estimated for the junction volumes N(0.0045, 0.0021) (in μm^3^); **b**. two thirds the estimated normal distribution for the junction volume; and **c**. half the estimated distribution for the junction volume. Thick solid black lines indicate the mean of the average edge traversal speed distribution reported in Ref. [11] (45.01 μm/s) and thick dotted black lines indicate mean ± standard deviation (45.01 ± 12.75 μm/s).

Further, the assumption that the entire junction volume is expelled during a contraction may not be realistic, as it would require extreme bending/extension of tubule walls and the ability of the entire junction to empty and fill out during each contraction. Reducing the volume Δ*V* expelled in each contraction to two-thirds of the estimated distribution of the junction’s volume (results shown in Fig. 7b) or to half (Fig. 7c) causes the average edge traversal speeds to drop considerably. The lower the proportion of the junction volume expelled during a contraction, the faster the pinches are required to be (i.e. large value of *α*) and the larger the frequencies of the pinch events (large *β*) in order to produce reasonably high average edge traversal speeds.

Next, we consider the particle transport generated by two types of ER sheets contracting and relaxing over a duration 2*T* (see Methods for details). The perinuclear sheets are shaped as contiguous layers of flat cisternae with a luminal volume larger than the tubules branching from these structures (see sketch in Fig. 1f). Accordingly, their contraction with 2*T* = 5 s and *V*_*sheet*_ = 10 μm^3^ yields a mean average traversal speed of 35 μm/s consistent with the single particle tracking experiments (see Fig. 8b). However, the speeds distribution tails towards higher values (Fig. 8a-b), something that has not be observed experimentally so far; it may be that high velocities cannot be recovered in experiments due to the constraints on linkage distance imposed to ensure trajectory fidelity in particle tracking.

**FIG 8:**
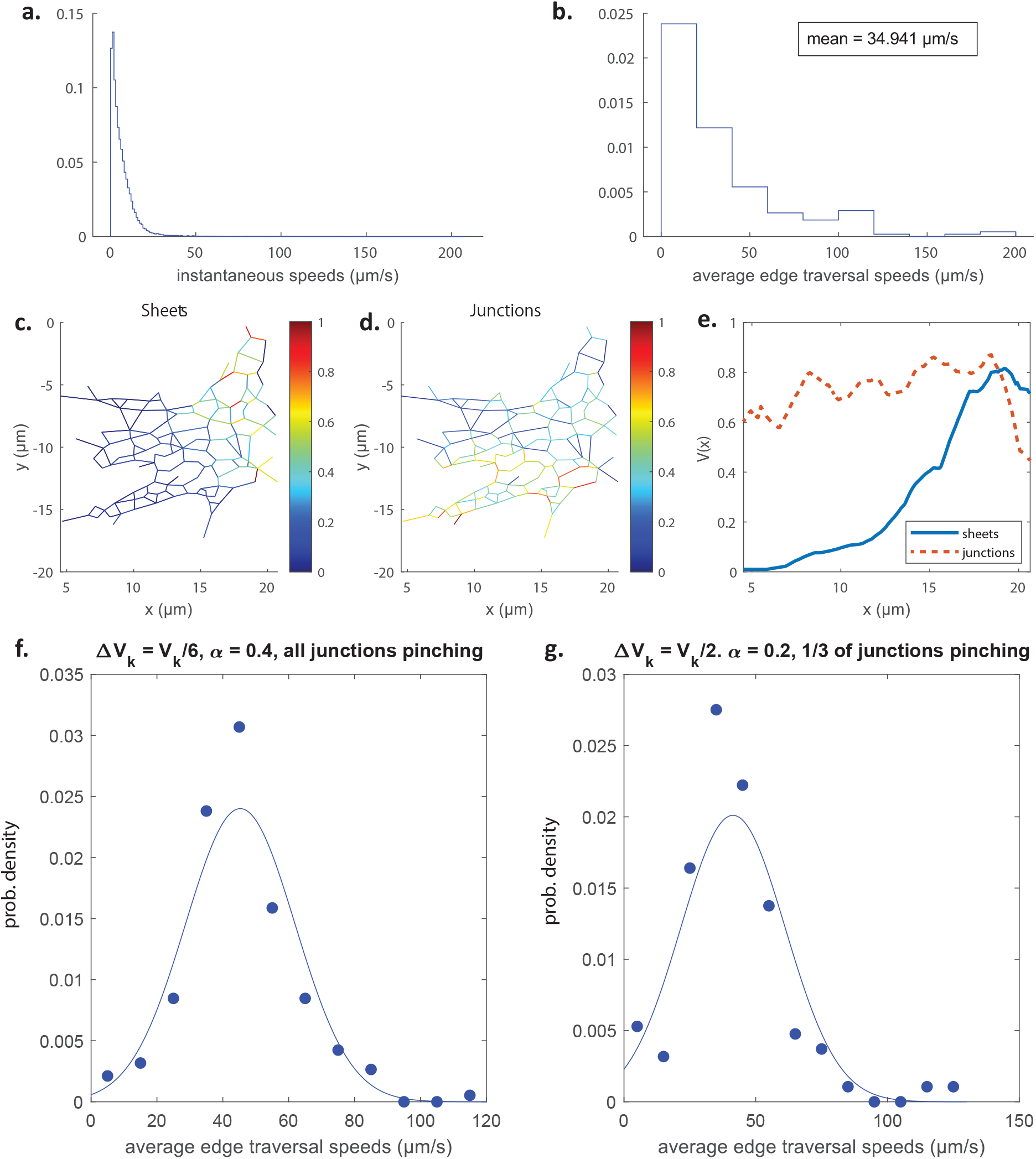
text on the next page FIG. 8: **a-b**. Distributions of instantaneous speeds (a), and average edge traversal speeds (b) obtained from simulations of the C1 network from Fig. 9 driven by the contraction of a perinuclear sheet. In these simulations the sheet undergoes one contraction+relaxation lasting 2*T* = 5 s, and expels a volume *V*_*sheet*_ = 10 μm^3^ of fluid during a contraction. **c-d**. Colour maps of normalised average edge traversal speeds obtained from simulations of the C1 network from Fig. 9 driven by contraction of tubules + sheets (c) and junctions + tubules (d) respectively. **e**. The speeds averaged along the *y* direction of the network, *V* (*x*), are plotted against *x*, to effectively project the information onto one dimension from c. (blue solid line) and d. (red dashed line). **f-g**. Histograms of average edge traversal speeds (dots) and normal fits (lines) and mean AETS (inset) in C1 networks with parameters adjusted as follows to approximate a network with peripheral sheets: Junction *k* expels a volume *V*_*k*_*/*6 of fluid in each pinch, the pinches are *α*^−1^ = 2.5 times slower than the original tubule pinches, and all nodes actively pinch (f); and node *k* expels a volume *V*_*k*_*/*2 of fluid in each pinch, the pinches are *α*^−1^ = 5 times slower than the original tubule pinches, and only a third of the nodes actively pinch (g).

Further, our simulations reveal that the flow decays sharply with distance away from the sheet where it originated as it branches out into tubules. This is illustrated in Fig. 8c-d and we further quantify the spatial gradient of the average edge traversal speeds across Cartesian 2D coordinates in Fig. 8e, revealing the stark contrast between the homogeneous profile for contracting junctions (red dotted line) vs sheet-driven transport (blue solid line). The short range of influence afforded by contracting perinuclear ER sheets thus argues against its ability to sustain mixing flows and fast particle transport on the distal tubular network.

Instead, we explore whether the peripheral sheets (i.e. the smaller flat inter-tubular ER regions, see sketch in Fig. 1d) may overcome the range limit. The peripheral sheets have volumes significantly larger than tubules and junctions, which we estimated at 0.12μm^3^ ± 0.04 μm^3^ (see Fig. 13 and Methods §IV J 4 for details). This suggests that the peripheral sheets could produce average edge traversal speeds compatible with experimental measurements. Assuming that a sheet typically occupies the area enclosed by a ‘triangle’ of tubules, we may incorporate the transport inside a sheet-driven network using our model for contracting junctions (also referred to as ‘nodes’ in our graph theoretical methodology), but with junction volumes set to the measured sheet volumes (see §IV J 4 for details) and with either (i) each node expelling one third of the volume expelled by a contracting sheet (since a peripheral sheet is in contact with three nodes on average), or (ii) a contracting node expelling the entire volume expelled by a contracting sheet, but with only one-third of the junctions actively contracting at any one time. Further assuming that during each contraction a peripheral sheet expels half its total volume so that in simulation (i) node *k* expels a volume *V*_*k*_*/*6 in each contraction and in simulation (ii) each active node *k* expels a volume *V*_*k*_*/*2, nodes which contract at rates *α*^−1^ = 2.5 and *α*^−1^ = 5 times slower than the tubule pinches in Ref. [11] in simulations (i) and (ii) respectively give average edge traversal speeds in the correct range of 40 μm/s (Fig. 8f-g). The contraction of peripheral sheets may thus be offered as a plausible mechanism for fast luminal transport, provided they are able to contract with sufficiently high rates.

Further, by relating the work done by a peripheral sheet contraction to the dissipation due to the flows induced inside the sheet and in the rest of the network, the energetic cost of one such contraction may be estimated to be of the order of 1000 molecules of ATP (see Methods §IV K for details). For reference, a kinesin motor protein uses one molecule of ATP to move 8 nm [17, 18], whereas a muscle fibre consumes hundreds of thousands of ATP per second [19]. These peripheral sheet contractions may be directly ATP-driven, or they might come as a result of other mechanical processes in the cell, similar to mechanisms generating cytoplasmic mixing flow [20–22].

## III. DISCUSSION

A better understanding of the interplay between ER structure and function via luminal transport speeds may hold clues to explaining the sensitivity of cells with extensive projections, such as sensory and motor neurons, to defects in ER shaping proteins. It is tempting to speculate that disturbed ER luminal transport, the kinetics of which is particularly important for communication across vast axonal lengths, underlies selective vulnerability of long neurons.

The motion of solutes in cellular compartments is now understood to be facilitated by active components. This is evident from direct motion measurements and the dependence of motion speed on the availability of ATP-contained energy [10, 11, 23, 24]. The origin of these active driving forces is, however, challenging to identify. The cytoplasm’s currents are often believed to originate from the motion of large complexes such as ribosomes and large vesicle cargo shuttled by cytoskeleton-mediated motorised transport [24]. In the case of enhanced transport in the lumen of the ER tubules, the contractility of tubules has been suggested as the flow generating mechanism, and indeed such tubule deformations have been observed in microscopy [11]. However, establishing a direct empirical link between tubule contraction and active flows, or experimentally testing other hypotheses for the driving mechanism behind ER solute transport, remain currently unattainable. In this study, we thus propose a physical modelling approach, which provides a platform to explore the nanofluidics behaviour of biological systems such as the ER network. The outcomes of our simulations for a contractile ER argue against the plausibility of local pinch-driven flow; pinches with frequency and size on the order of those estimated by microscopy yield significantly lower speeds than single particle tracking measurements, as well as no enhancement of mixing beyond that from passive diffusion. The deficit stems from the fact that the displaced fluid volume upon local contraction is too small to generate sufficient particle transport.

Given the uncertainty of the empirical measurements for ER tubule deformation, due to the limits in the spatio-temporal resolution of organelle structure imaging, the pinch parameters may have been significantly underestimated. This sanctioned exploring a wider range of spatio-temporal parameters in our simulations, which revealed that a combination of a higher frequency with a much larger pinch length may provide higher particle speeds that are comparable to the single particle tracking measurements. Further, our modelling results suggest the possibility of transport by luminal width contractions of larger volume ER subdomains (which are contiguously interconnected with the network). In that respect, the contraction of peripheral (i.e. inter-tubular) sheets, in particular yields speed values in a plausible range provided they are able to contrast fast enough. In contrast, the contraction of large-volume perinuclear sheets leads to fast transport but with a limited spatial range and with flows that would not impact transport beyond a few microns into the peripheral network. Similarly, the alternative scenario of contractility of tubular junctions appears unlikely as particle speeds similar to experiments could only appear for junctions pinching at up to 8 times faster than experimentally observed contracting tubules.

The modelling approach in this study, although focused on the ER network, provides a step forward towards understanding intra-organellar fluid dynamics. Our simulation results have been used to rule out several scenarios, which seemed physically intuitive but are nevertheless unable to explain the observed enhanced luminal transport. Furthermore, our results generated a set of potentially testable predictions that can be used to validate or refute each of the envisaged transport mechanisms. For example, as fluid expulsion from peripheral sheets appears to be in broad in agreement with single particle tracking measurements data under even conservative assumptions, future measurements may explore whether an active luminal motion is more pronounced and faster in proximity to peripheral sheets. Moreover, a set of improved spatio-temporal resolution measurements of particle tracking and structural contractions will be needed to complete the physical picture of ER luminal transport.

While our study allowed us to test different plausible scenarios, questions remain open as to the force generation mechanisms responsible for the observed pinching dynamics. We have estimated in §IV H that forces on the order of 30 pN would be required to periodically contract the tubules as seen empirically. The force exerted by a single molecular motor has been estimated to be on the order of 6 pN [25], so the required forces for pinching may be provided by several motors working together [26]. Recently discovered hydro-osmotic instabilities could also contribute to shape fluctuations and pinching, although they are predicted to have much longer wavelengths than the typical size of a pinching region [27]. In contrast, the mechanisms involving topological remodelling of the ER, such as the well-documented process of ring closure [28], occur over timescales of minutes and therefore cannot account for the millisecond-scale transport measured here.

We have used *in silico* fluidics modelling for our conclusions, particularly in identifying new physically permitted mechanisms of luminal propulsion, and our simulations should be regarded as theoretical predictions demonstrating physical plausibility rather than mere speculation. We have explored theoretically the consequences of structural fluctuations which presumably do take place. Fluctuations in width of the flat ER areas, in particular, are expected since the structures appear dynamic in live light microscopy and narrowing/extension points are observable in electron microscopy [29, 30]. Further, variability in the areas of flat ER that are observable in electron microscopy can be explained by capturing the structures in different states of fluctuations. It should be expected that the mobile elastic structure such as ER sheets would not exhibit rigidity required to prohibit fluctuations.

In conclusion, it is worth emphasising that our *in silico* fluid dynamical modelling reveals that for structural fluctuation-based mechanisms to facilitate luminal motion, assumptions currently not supported by empirical data are required. This warrants explorations of alternative mechanisms for ER luminal transport, for example anomalous diffusion driven by the fluctuations of macromolecular complexes [24] or by osmotic forces, as previously suggested [11].

## IV. METHODS

Here we describe the mathematical and physical model for the fluid mechanics and transport driven by active contractions in the ER. This requires the introduction of a network model for the geometry of the ER (§IV A) and individual pinches (§IV B) and a framework for the hydrodynamics of pinching tubules (§IV C). Our solution method for the flows inside our network is then described in §IV D. From simulations of Brownian particles advected by these flows (§IV E) quantitative measures of particle transport (§IV F) are extracted for comparison with experiment. Boundary slip is incorporated into our model in §IV G. The force required to pinch a tubule is estimated in §IV H. In §IV I we present in detail the derivations of the theoretical results discussed in §II C (transport upper bound by a single pinch) and §II D (coordination of pinches). Finally, modifications of our model to explore alternative flow generation mechanisms are discussed in §IV J, and the energetic cost in contracting a peripheral sheet is estimated in §IV K.

### A. Network modelling

We represent the ‘skeleton’ of a two-dimensional ER network as a planar graph with each node assigned a position **x** ∈ ℝ ^2^. Given an edge of the network labelled (*i, j*) and of length |**x**_*i*_ − **x**_*j*_| = *l*, we model the lumen of the corresponding tubule to occupy a cylinder of radius *R* whose axis lies along the edge and has length *l*. This assumption avoids the intrinsic difficulty in defining a precise boundary between a tubule and a tubular junction, as well as leading to a simplified model of the intra-nodal dynamics of a solute particle (see below); since the tubules are long compared to the size of the junctions, the impact of their small overlap can be safely neglected.

We mathematically reconstruct a model ER network, which we refer to as C0, using the skeleton image of the COS7 ER network given in Fig. 4 of the Supplementary Material from Ref. [11]; this network is reproduced in Fig. 9a. We use the multi-point tool in ImageJ [31] to place numbered points at the positions of the nodes on the source skeleton image, and then obtain a list of the indices and position coordinates of each node. The edges are then manually tabulated as a list of pairs of nodes; this gives us all the information required to construct a mathematical graph as shown in Fig. 9b, with the original network superimposed on the mathematical model in Fig. 9c. Note that in what follows we work with the largest connected component of the source skeleton image in order to study transport in a fully connected network. We also use the same procedure to extract the graph structures from microscopy images, of four smaller ER networks which we label C1-C4 (original network with mathematical graph superimposed in Fig. 9d). In Fig. 9e, we show the distributions of the edge (tubule) lengths in each of the C0-C4 networks, as well as the mean edge lengths, the mean degrees (the degree of a node is the number of edges connected to it), and the number of nodes. The mean edge lengths are around 1 μm and mean degrees are approximately 3. In order to compare the biological network to an idealised ER system, we also consider a honeycomb network, i.e. one where every node (apart from those at the boundaries) has a degree of 3, with all edge lengths exactly 1 μm.

**FIG 9:**
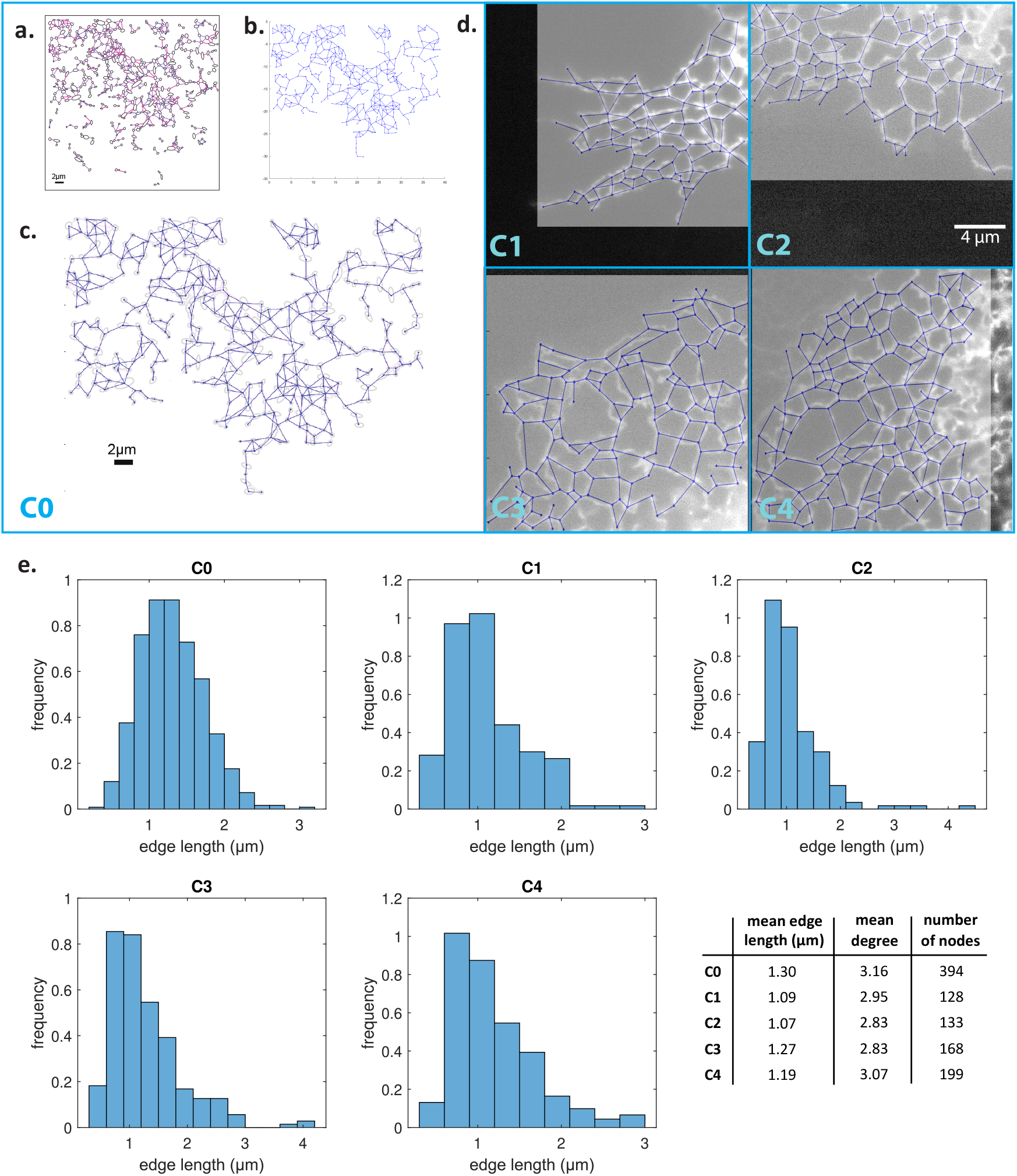
**a**. Skeleton image of COS7 ER reproduced from Ref. [11]. **b**. Model ER graph (blue solid lines) reconstructed from (a) using ImageJ. **c**. Experimental images in (a) superimposed with mathematical model from (b). **d**. Microscopy images of four different COS7 ER networks (labelled C1-C4) with reconstructed model networks (blue solid lines) superimposed. **e**. Distributions of edge lengths in the C0-C4 networks. Bottom right: mean edge lengths, mean degrees (i.e. number of edges connected to a node) and number of nodes of the C0-C4 networks.

### B. Pinch modelling

To describe the pinches (Fig. 1e), we consider a model where the kinematics of each pinch is fully prescribed in time. We therefore assume that the active biochemical forces responsible for the deformation of the tubules balance with the elastic resistance of the tubules and with the dissipative forces in the fluid in such a way that the pinches occur as described.

The geometrical model for a pinch is illustrated in Fig. 10. Each tubule is assumed to have a pinch site at its midpoint (i.e. with *L*_1_ = *L*_2_); we make this simplifying assumption in our simulations, after verifying that more general pinch locations (*L*_1_ ≠ *L*_2_) have virtually no effect on particle transport. The pinching events occur at the pinch sites stochastically and independently of each other. Each pinch is defined geometrically by three parameters from a random distribution (see below): (i) the duration of a pinch 2*T*, (ii) the time *T*_*wait*_ between the end of a pinch and the beginning of a new one on the same site, and (iii) the length of a pinch 2*L* (see Fig. 10). We assume for simplicity that all pinches are axisymmetric so that using the notation of Fig. 10, each tubule remains a cylinder of time-varying radius denoted by *r* = *a*(*z, t*). Pinches are assumed to have reflectional symmetry about the plane in the cross-section through the centre of the pinch (i.e. Fig. 10 each pinch is characterised by the same length *L* on either side of it). The total length of a tubule is denoted by *l* = *L*_1_ + 2*L* + *L*_2_, so the portion before the pinch has length *L*_1_, and that after the pinch is of length *L*_2_.

**FIG 10:**
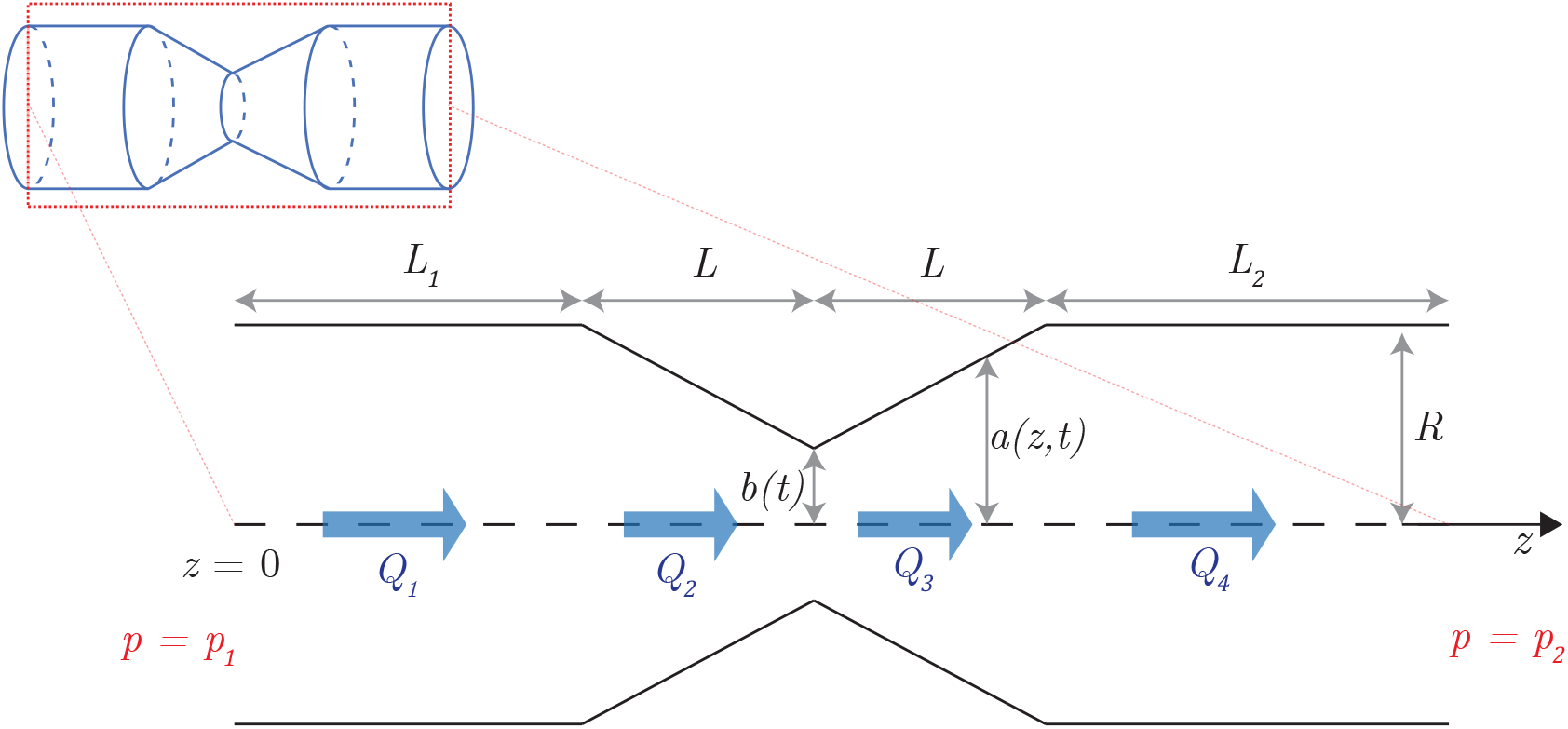
Mathematical model of a pinching tubule. The tubule has a radius of *R* outside the pinch, *a*(*z, t*) in the pinch (where *z* is the axial coordinate), and *b*(*t*) at its narrowest point i.e. the centre of the pinch. The portion of the tubule before the tubule has length *L*_1_ while that after has length *L*_2_; the pinch is symmetric and has a length 2*L. Q*_1_, *Q*_2_, *Q*_3_, and *Q*_4_ denote the volume fluxes through the tubule in the four different regions as indicated. The pressures at the end of the tubule are *p* = *p*_1_ at *z* = 0 and *p* = *p*_2_ at *z* = *L*_1_ + 2*L* + *L*_2_.

We model the geometrical profile of each pinch of length 2*L* as following a linear radius change (see Fig. 10). Within a pinch located at *z* = *z*_0_, the radius of the cylinder is therefore given by

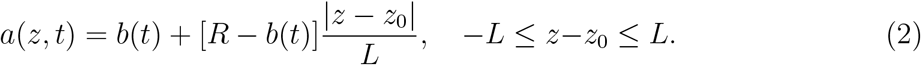

The time-varying function *b*(*t*) ≤ *R* is therefore the minimum pinch radius in the centre of the tubule. As the simplest modelling choice, we assume in our simulations that *b* changes in time sinusoidally and thus will use *b*(*t*) = (*R* + *b*_0_)*/*2 + (*R* − *b*_0_) cos(*πt/T*)*/*2, where *t* is time after the pinch begins and *b*_0_ is the value of *b* halfway in time through the pinch. Choosing a smooth time variation for the function *b*(*t*) will ensure the continuity of fluxes in time (see Eq. (10) later). We have verified that changing the pinch shape to a smoother geometry has essentially no impact on the fluid volume expelled/taken in, and therefore no significant effect on the flows/transport.

In our simulations the stochastic pinch parameters are drawn from the distributions measured in Ref. [11]. The pinch duration 2*T* is therefore drawn from an exponential distribution with rate parameter *λ* = ln 10*/*0.167 s^−1^ (so the mean value is 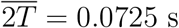), the time between pinches *T*_*wait*_ from from an exponential distribution with rate parameter *λ*_*wait*_ = ln 10*/*0.851 s^−1^ (mean value 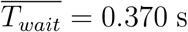) and the pinch length 2*L* from a uniform distribution with mean *μ* = 0.12 μm and variance *σ*^2^ = (0.06^2^*/* ln 10) μm^2^ ≈ (0.040 μm)^2^. Throughout we set the tubule radius *R* = 30 nm and *b*_0_ = 0.01*R* (recall from Fig. 10).

### C. Hydrodynamic modelling

#### 1. Hydrodynamics of network

We assume that the fluid occupying the ER network is Newtonian. Flow inside the network occurs at low Reynolds number, which can be justified as follows. From experiments in Ref. [11], we know that velocity scale relevant to ER flows is of the order *U* ∼ 10^−5^ ms^−1^. With a typical tubule radius *R* ≈ 10^−8^ m and a kinematic viscosity of at least that of water *ν* ≈ 10^−6^ m^2^ s^−1^, we obtain a Reynolds number of the order Re = *UR/ν* ∼ 10^−7^, so the flow is indeed Stokesian and inertial effects in the fluid can be safely neglected.

At a typical instance in time, a tubule undergoing contraction (or relaxation) causes a net volume flux to exit (or enter) the corresponding tubule. In the context of our graph theoretical model, we therefore model each pinch site as a ‘pinch node’ generating a net hydrodynamic source/sink whenever the tubule contracts/relaxes.

However, since it is not guaranteed that the total volume created by all pinching events always adds up to zero, we need a mechanism for the corresponding net volume to exit, or enter, the network. We achieve this through a number of ‘exit nodes’ that allow mass to be globally conserved. We locate the exit nodes at the exterior of the network and choose their number randomly. From a hydrodynamic standpoint, we impose the pressure condition *p* = *p*_0_ at each exit node to model their connection to a large fluid reservoir.

We numerically tested the robustness of our results to the details of the exit nodes by repeating simulations with different configurations. The exact choices of exit nodes turn out to not affect the transport results shown below provided there are sufficiently many of them to avoid channeling the entire network’s worth of pinch-induced flow into a few tubules towards the exterior, thereby producing artificially strong flows.

#### 2. Hydrodynamic model for a pinch

### a. Flow rate

The velocity field in a straight cylindrical tubule at low Reynolds number is the classical parabolic Poiseuille flow [32]. When integrated over a cross section of the tubule, this flow yields the Hagen-Poiseuille law relating the pressure change Δ*p* across a length *l* of a tubule to the a net volume flux *Q* in the positive axial direction,

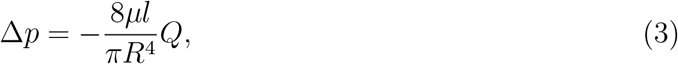

where *μ* is the dynamic viscosity of the Newtonian fluid. Note that when *Q >* 0 our notation leads to Δ*p <* 0, meaning that the pressure decreases across the length of the channel.

The result in Eq. (3) is valid for a straight (i.e. not pinching) tubule, and we need to generalise it to the case of a pinching tubule. Consider first a more general axisymmetric pipe whose radius *a*(*z, t*) varies with axial position *z* and time. We may use the long-wavelength (lubrication) solution to Stokes’ equations for the streamwise velocity *u*(*z, r, t*) and flux *Q*(*z, t*) inside such a pipe into which a flux *Q*_1_(*t*) enters at *z* = 0 [33],

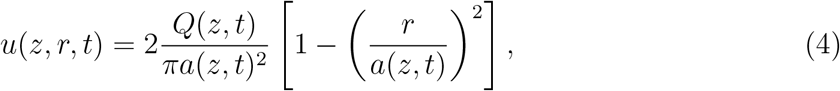

Where

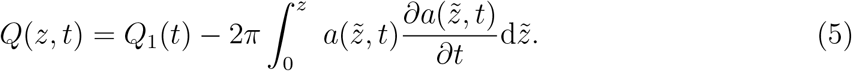

The equality in Eq. (5) can be derived using an intuitive mass conservation argument, independently of the inspired ansatz in Eq. (4). Conservation of mass inside a small section [*z, z* + *δz*] of the cylinder requires 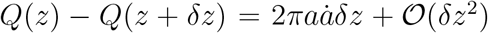. Considering the limit *δz* → 0 and integrating the resulting expression for *∂Q/∂z* from 0 to *z* yields Eq. (5).

In order for the no-slip boundary conditions at *r* = *a*(*z, t*) to be satisfied, and also to satisfy the incompressibility condition ∇ · **u** = 0, the radial component of the velocity, *v*(*z, r, t*), is necessarily non-zero and given by [33]

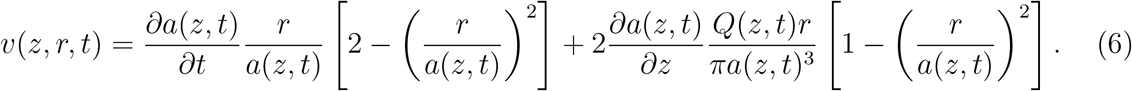

Our geometrical model for each pinch in §IV B yields straightforwardly a piecewise linear expression for *a*(*z, t*) in different regions of the tubule, with the time dependence entering only through the value of the pinch radius *b*(*t*). Denoting by *Q*_*i*_ (1 ≤ *i* ≤ 4) the fluxes in the four regions of the pinched tubules shown in Fig. 10, we may substitute the linear shape functions into Eq. (5) and obtain

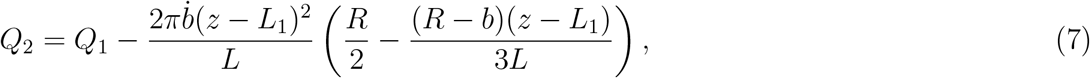

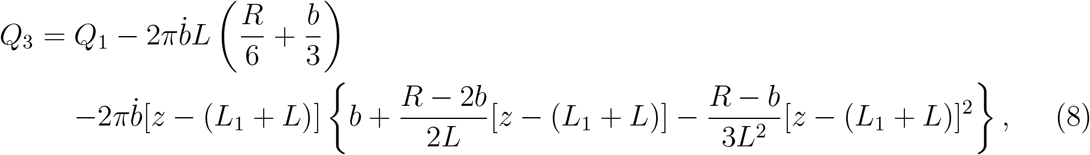

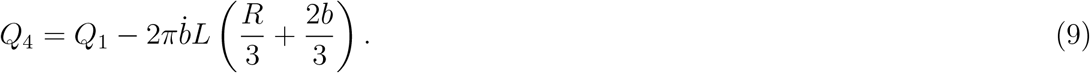

Note that the final expression may be rearranged as

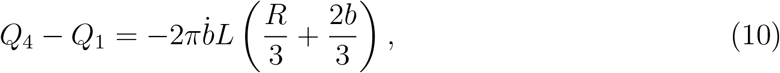

which may be interpreted as the instantaneous volume source/sink during a contraction/relaxation at a pinch site.

### b. Pressure drop

We next need to compute the pressure drop in the pinches. We integrate the *z*-component of the Stokes equation

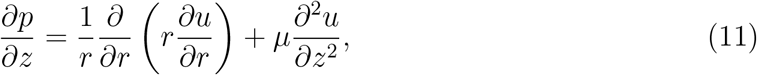

along 0 ≤ *z* ≤ *L*_1_ + 2*L* + *L*_2_, and use the solution for *u*, to obtain

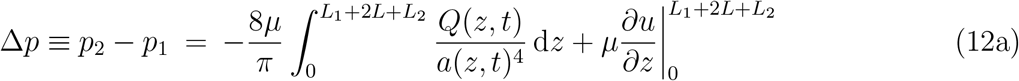

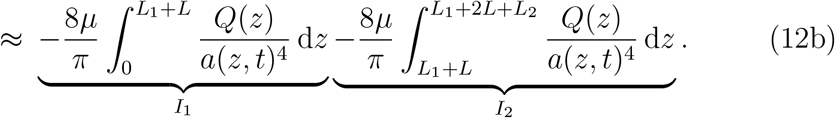

Here, the second term on the right-hand side of Eq. (12a) has vanished because *∂*_*z*_*u* ∝ *∂*_*z*_(*Q/a*^2^) but *Q* and *a* are approximately constant at the entrance and exits of the tubule when the pinch site is sufficiently far from the ends of the tubule so that the flow is fully-developed there.

Using the expression (7) for *Q*_2_, an integration yields

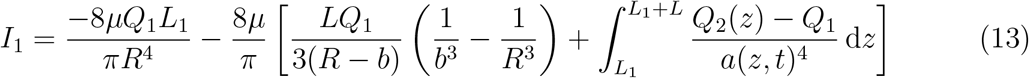

and, by symmetry,

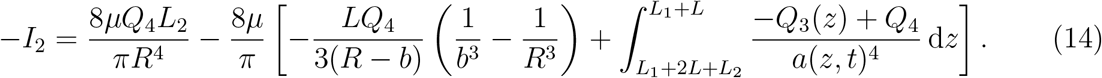

Subtracting these two results (and noting the integrals cancel out by symmetry) we obtain the modified Hagen-Poiseuille expression for a pinching tubule as

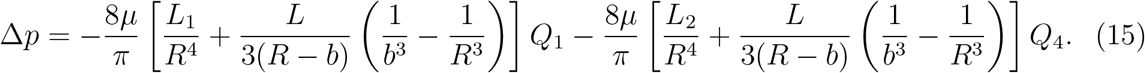

Note that this relationship is linear in each of *Q*_1_ and *Q*_4_, and is to be solved alongside Eq. (10) to relate the flow rates and pressure drops to the change in size of the pinches. Importantly, the classical Hagen-Poiseuille law is recovered as *b* → *R* since Eq. (15) becomes in that limit

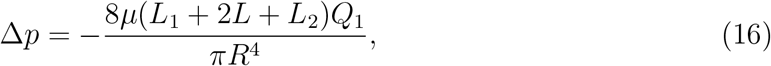

which agrees with Eq. (3) when taking *l* = *L*_1_ + 2*L* + *L*_2_.

### D. Solving the hydrodynamic network model

The incorporation of pinches as ‘dummy nodes’ into the graph theoretical framework of §IV A along with Eqs. (10) and (15) for the necessary pinch-related quantities allow us to reduce the problem of determining the time-dependent flows in an active pinching network into the simpler problem of solving at each instant for the instantaneous fluxes inside a ‘passive’ network with newly added nodes, appropriate sources/sinks, and modified pressure drops. Note that since the flows at these sub-cellular scales are inertialess (i.e. Stokes flows), we are able to effectively decouple time from our problem and solve the problem in the quasi-steady limit.

For each edge (i.e. tubule) (*i, j*) in the network, we define *Q*_*ij*_ to be the flow rate from node *i* to node *j*, with the sign convention that flow is from *i* to *j* if *Q*_*ij*_ *>* 0. For mathematical convenience, we define *Q*_*ij*_ = 0 in all cases where (*i, j*) is not an edge in the graph. The goal is to solve for the values of the *Q*_*ij*_’s corresponding to each edge.

After the incorporation of the dummy nodes, we denote by *N* the number of nodes and *E* the number of edges. We label the nodes such that {1, …, *M* } denotes the *M* exit nodes. Let *q*_*i*_ be the source or sink carried by the *i*^th^ node (so that *q*_*i*_ = 0 if *i* is a normal node, *q*_*i*_ is as specified by the RHS of Eq. (10) if *i* is a pinch node, and *q*_*i*_ is a quantity to be determined if *i* is an exit node.). Our *M* + *E* independent variables are therefore {*q*_*i*_|*i* = 1, …, *M* } i.e. the sources/sinks carried by the exit nodes, and the *Q*_*ij*_’s corresponding to each edge. To obtain their values, we employ the viscous hydraulic analogues of Kirchoff’s Laws.

#### 1. Kirchhoff’s First Law (K1)

The first equation is that mass is conserved at each junction i.e. for each node *i* we have

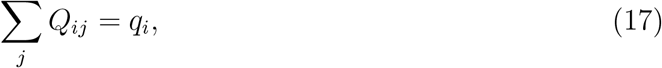

which gives us therefore *N* equations. Note that these equations together imply global conservation of mass,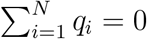.

#### 2. Kirchhoff’s Second Law (K2)

The second equation is a statement of consistency of pressure, namely that the pressure change around any cycle (i.e. closed loop) of the network is zero. Therefore, in a given cycle *C* = {*v*_1_, *v*_2_, …, *v*_*n*_, *v*_*n*+1_ = *v*_1_}, if 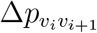 denotes the pressure change from node *v*_*i*_ to node *v*_*i*+1_ we necessarily have

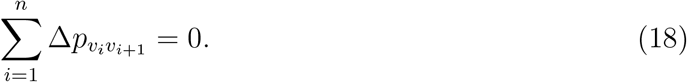

Note that the pressure change across a node is negligible.

The K2 statement in Eq. (18) applies to all cycles in the graph, which would give us more equations than we need since the vectors of coefficients in Eq. (18) are linearly dependent. Instead, we need a minimal set of linearly independent K2 equations, corresponding to the cycles in a cycle basis of the graph, and we need only apply K2 to these cycles.

To construct a cycle basis of the graph *G*, we use standard results from graph theory [34]. We first construct a spanning tree *T*, defined as a connected subgraph which contains all the nodes of *G* and no cycle, as shown in Fig. 11a on an example. Any tree *T* has *N* − 1 edges. Therefore there are *E* − (*N* − 1) edges in the graph *G* but not in the spanning tree *T* ; for each such edge *e*, we denote by *C*_*e*_ the unique cycle in the graph created by adding the edge *e* to the tree *T* (see Fig. 11b). The set of all such cycles *C*_*e*_ is then a cycle basis of *G*, and thus there are *E* − (*N* − 1) cycles in this set. There are therefore *E* − *N* + 1 independent cycles in the cycle basis.

**FIG 11:**
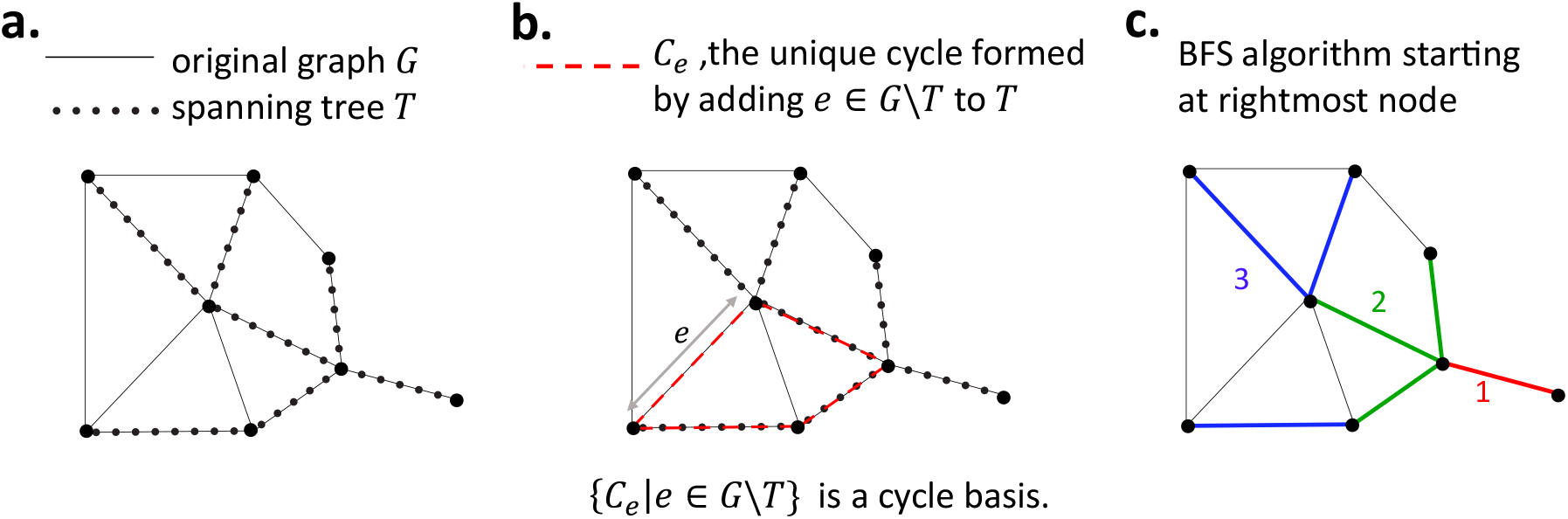
Elements of graph theory required to model the ER network. **a**. A graph *G* (black solid lines) and its spanning tree *T* (black dots). **b**. The unique cycle *C*_*e*_ (red) formed by adding an edge *e* ∈ *G*\*T* to *T*. **c**. A breadth-first search (BFS) starting at the rightmost node; the graph is explored in the order red, green, blue.

To compute the spanning tree *T* we use a breadth-first search (BFS) algorithm [35]. We start from an arbitrary node and explore its neighbouring nodes. We add to *T* any previously unexplored node (and the corresponding edge) which does not result in the creation of a cycle in *T*. We then repeat this (in an arbitrary order) on the neighbours of the previous generation of nodes added to *T*, until no more nodes are left to explore. This is illustrated on an example in Fig. 11c.

A similar algorithm is also used to compute the cycles *C*_*e*_ in the cycle basis. This time, however, denoting *e* = (*i, j*), the algorithm is started from *i* and set to terminate as soon as *j* is visited, yielding a path in the tree from *i* to *j*, which, together with the original edge (*i, j*), completes a cycle.

#### 3. Pressure boundary conditions at exit nodes

At this point in the modelling, we have *M* + *E* independent variables (the flow rates in each edge and at the exit nodes), *N* equations from K1 (i.e. Eq. (17)), and *E* − *N* + 1 equations from K2 (i.e. Eq. (18)). The remaining *M* − 1 equations follow from the pressure conditions at the exit nodes. Specifically, the pressure difference between any pair of exit nodes *i* and *j* is zero

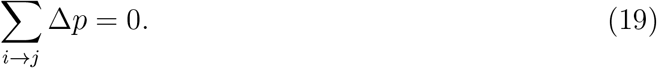

Here the sum Σ_*i*→*j*_ is defined to be over any path *P*_ij_ from *i* to *j*. Thanks to Kirchhoff’s second law, this quantity is path-independent. This indeed gives us 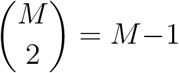 equations, which we pick to be Σ _1→*j*_ Δ*p* = 0 for *j* = 2, …, *M*.

We then solve the resulting linear system of *M* + *E* equations numerically. Note that we do not need to specify the value of the fluid viscosity *μ* in our algorithm because it cancels out in the K2 equation, Eq. (18), and in the pressure boundary condition equation, Eq. (19).

### E. Simulating particle transport

With our solution for the flows in the active network at each instant of time, we now proceed to track the motion of Brownian particles inside the network using a discretisation of their stochastic equations of motion, as a model for the transport of proteins in the ER network.

We use the simplest approach where we superimpose Brownian motion onto advection by the flow inside each tubule. Let **x**(*t*) denote the position of a Brownian particle in a tubule and **x**_*n*_ the finite-difference approximation of **x**(*n*Δ*t*), where Δ*t* is a discrete time step. The displacement of the particle at each time step can be obtained approximately using an explicit first-order Euler scheme

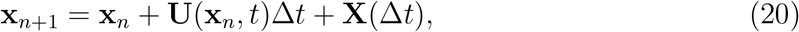

where **U** is the instantaneous flow velocity, and the random noise term **X**(Δ*t*) is drawn from a zero-mean Gaussian with variance ⟨**X**(Δ*t*)**X**(Δ*t*)⟩ = 2*D*Δ*t***I**, where *D* is the Brownian diffusivity of the particle. In our simulations, we take the diffusion constant to be the mean intranode diffusivity measured in Ref. [11], *D* ≈ 0.6 μm^2^s^−1^. We include interactions between particles and walls by assuming that particles perfectly reflect off walls (i.e. elastic collisions). The particles are modelled as rigid spheres of diameter 5 nm, and the size of the particle matters only during elastic collisions with the walls of the tubules. As relevant in the limit of low volume fraction, we neglect hydrodynamic interactions between particles and perform ensemble averaging of the trajectories of many independent particles.

When a particle enters a node, we model its dynamics as follows. We consider a particle to have entered a node only if it has reached the end of a tubule, say of length *l*, at which instance we assign the particle to the node-point (i.e. the single point associated with the node in the graph description of the ER network). Although nodes contain a three-dimensional volume, their typical nodal length scale is of the order *R* ≪ *l*, and thus approximating them by point nodes is appropriate on the scale of the whole network. To decide towards which of the connected tubules the particle leaves the node, we estimate the values of the Péclet number Pe in each of the tubules. We define a local Péclet Pe_*i*_ = *U*_*i*_*R/D* where *U*_*i*_ is the mean flow velocity through tubule *i*, with *U*_*i*_ *>* 0 for flow out of the node and *U*_*i*_ ≤ 0 otherwise. We then assume that the particle enters a neighbouring tubule *i* connected to the node with a probability proportional to max(Pe_*i*_ + 1, 0). This ensures that we have the expected behaviour in both limits of Pe: at high (positive) Péclet numbers, the probability is proportional to the flow speeds in each of the connected tubules, while at low Péclet the exit of the node is limited by diffusion and thus the exit is equally likely in each tubule.

### F. Data processing: instantaneous speeds and average edge traversal speeds

During each simulation, we compute the edge traversal speeds as follows. A particle is defined to traverse an edge (*i, j*) if it travels from node (i.e. junction) *i* to node *j*, or from *j* to *i*, without visiting *i* or *j* in between. The corresponding edge traversal time is then the time between the arrival at the target node and the most recent departure from the node of origin. The edge traversal speed is naturally defined as the length of the tubule (*i, j*) divided by the edge traversal time.

The average edge traversal speed associated with an edge (*i, j*) is then defined as the mean over all edge traversal events across (*i, j*) of the edge traversal speeds.

In addition we also compute for each particle the ‘instantaneous’ speeds defined by *V*_*n*_ = |**X**(*t*_*n*+1_) − **X**(*t*_*n*_)|*/*Δ*t*, where *t*_*n*_ = *n*Δ*t* with Δ*t* = 18 ms, which is the same temporal resolution as in the particle tracking carried out in Ref. [11].

### G. Incorporating slip boundary conditions

The methodology we have detailed thus far assumes no-slip boundary conditions at the tubule walls for the fluid flow. However, the membrane-bound lipids themselves could also flow in response to the nanoscale luminal flows. This may be modelled by introducing a finite slip boundary condition on the tubule wall. The slip boundary conditions with a slip length *λ* ≥ 0 at the wall *r* = *a*(*z, t*) are given by

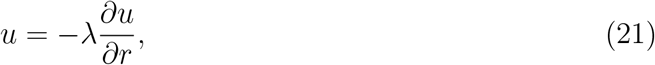

and 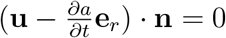, with 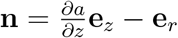, which simplifies to

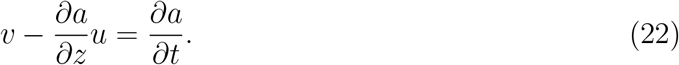

We may then derive the long-wavelength solution for the flow field inside an axisymmetric deforming tubule as follows. Using an ansatz for the axial component *u* that is motivated by the uniform-radius Poiseuille flow with slip,

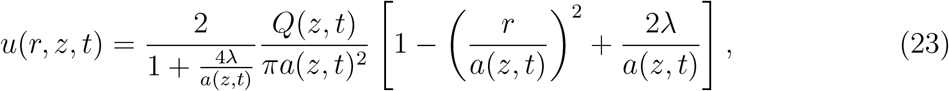

we may solve the incompressibility condition 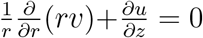. Regularity at *r* = 0 constrains the integration constant to be zero, yielding

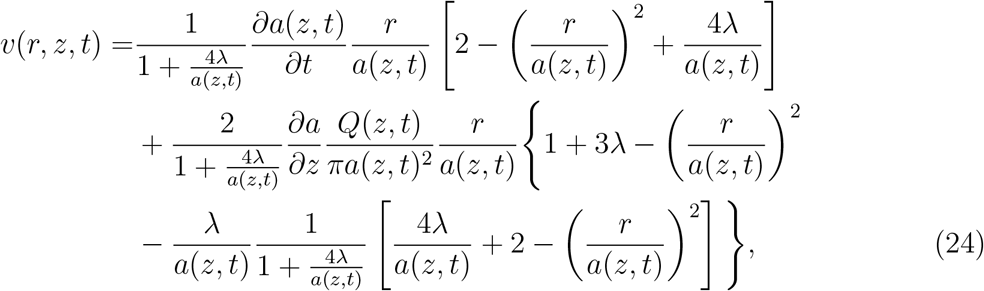

which automatically satisfies the boundary condition in Eq. (22). Note that the mass conservation equations are not affected by the introduction of a slip length.

Using the new solution for *u*, the modified Hagen-Poiseuille expression with slip may be derived as before to be

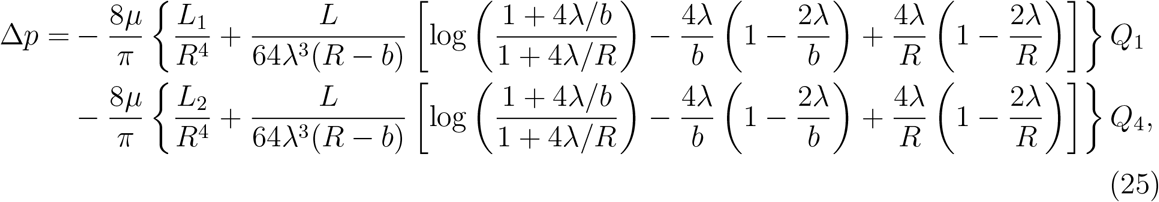

which does recover the no-slip result as *λ* → 0.

These results may then be used to simulate particle transport with slip boundary conditions. In Fig. 3a we plot the distributions of average edge traveral speeds obtained from simulations of a C1 network pinching with the original pinch parameters from Ref. [11] for four different values of the boundary slip length. In Fig. 3b we further display the profiles of the longitudinal flow (Eq. (23)) for different slip lengths with the volume flux fixed.

### H. Estimate of forces required for pinches

In this section we derive an order-of-magnitude estimate for the forces required to pinch an ER tubule.

We first estimate the difference in the membrane’s elastic energy, Δ*E* = *E*_pinched_ − *E*_unpinched_, between the pinched (*E*_pinched_) and unpinched configurations (*E*_unpinched_). In the absence of spontaneous curvature, the Helfrich free energy *h* per unit area of a membrane is given by

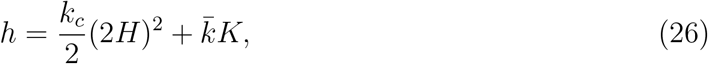

where *k*_*c*_ and 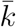 are bending rigidities, *H* is the mean curvature, and *K* is the Gaussian curvature [36]. The mean and Gaussian curvatures may be expressed in terms of the principal curvatures *κ*_1_, *κ*_2_ as *H* = (*κ*_1_ + *κ*_2_)*/*2 and *K* = *κ*_1_*κ*_2_. We take *κ*_1_ and *κ*_2_ to be the principal curvatures in the directions normal and parallel respectively to the tubule’s longitudinal axis.

The dominant contribution to Δ*E* is from *E*_pinched_, specifically from the region near the pinch site i.e. the centre of the pinch, where the tubule radius is smallest, taken to be at *z* = 0. We have *κ*_1_ = 1*/a*(*z, t*) and *κ*_2_ = 𝒪 (*R/L*^2^), yielding *H* = 1*/*2*a* + 𝒪 (*R/L*^2^) and *K* = 𝒪 (*R/aL*^2^). The dominant contribution to the membrane energy 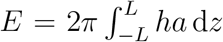 is therefore from *H*, in a neighbourhood of *z* = 0. Since real pinches do not have kinks, a linear term, for the purposes of estimating membrane energy, is unphysical, and we expect *a*(*z, t*_0_) = *b*_0_ + 𝒪(*z*^2^) near *z* = 0, where *b*_0_ is the pinch radius in the centre of the pinch in the maximally contracted state and *t*_0_ denotes a time at which the tubule is in a pinched state. We may therefore write *a*(*z*) = *b*_0_ + *Rz*^2^*/L*^2^ to obtain an order-of-magnitude estimate of the membrane energy in the pinched state.

Evaluating the integral for *E*_pinched_ then yields the leading-order estimate

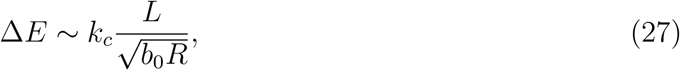

We may take 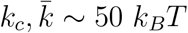(*k*_*B*_ is the Boltzmann constant times and *T* the room temperature) [37], with estimate values *R* = 30 nm, and *L* = 70 nm. The minimum pinch radius *b*_0_ may be estimated to be 10 nm allowing for membrane thickness and incomplete squeeze, yielding

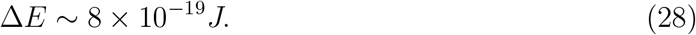

The hydrodynamic contribution to the energy expenditure during a pinch may be calculated as the sum of the dissipation inside the pinch and in the rest of the network (see §IV K for an analogous calculation for a contracting peripheral sheet). We find that the hydrodynamic contribution is negligible compared to the elastic component of the work done to pinch a tubule.

An estimate for the force required to pinch the tubule may then finally be obtained as *F* ∼ Δ*E/R* ∼ 30 pN.

### I. Derivation of theoretical bounds for active flows driven by pinching tubules

#### 1. Advection due to a single pinch

In this section we calculate an upper bound for the axial distance Δ*z* a particle can be advected by the flow produced by an individual pinch. Recall the formula for the volume ‘source’ due to a pinch,

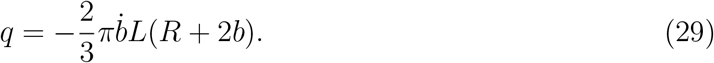

Among all possible ways for a particle to be transported by a pinch, an upper bound on the transport distance Δ*z* ≤ Δ*z*_max_ can be reached if all of the following conditions are satisfied: (i) All of the source flows to one side of the pinch (i.e. there is no leakage on the other side); (ii) The particle travels outside the pinching region (axial flows within the pinching region produce smaller advective displacements than those outside, as may be verified numerically); (iii) The particle travels along the centreline of the tubule (i.e. at twice the cross-section averaged flow velocity, a standard Poiseuille result); (iv) The minimum pinch radius *b*_0_ (recall from Fig. 10) is 0.

Under conditions (i) and (ii), the cross-section averaged speed corresponding to maximal transport generated from a single pinch can then be computed using Eq. (29) as

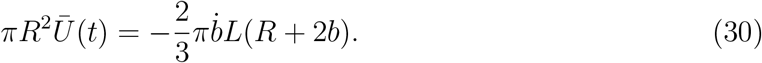

Using condition (iii), the position of the particle along the centreline *z*(*t*) satisfies then the ordinary differential equation

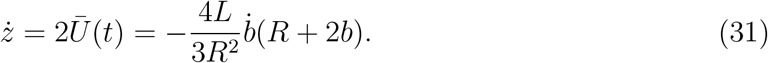

Integrating this equation from *t* = 0 (start of contraction, with *b*(0) = *R*) to *T* (end of contraction, with *b*(*T*) = *b*_0_) then leads to

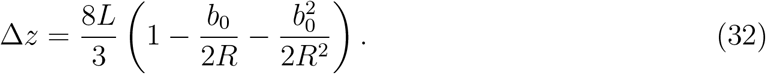

The value of Δ*z* is maximized when *b*_0_ = 0 (i.e. when condition (iv) holds), yielding the upper bound

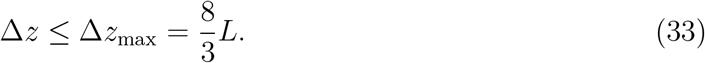

#### 2. Extension to nonlinear interactions between two pinches

An isolated pinch is only capable of reciprocal motions. The simplest system capable of producing non-reciprocal motions is illustrated in Fig. 5 and consists of two pinch sites arranged in series near the midpoint of a long horizontal tubule.

We may calculate an upper bound on the net particle displacement that can be achieved after both pinch sites pinch exactly once. We make the assumptions (ii)-(iv) as above, and in addition, allow pinches to spend extended amounts of time in their completely closed state; any deviation from these assumptions will result in net transport that is further reduced.

How can we maximise the positive (i.e. rightward in Fig. 5) particle displacements induced by the contractions, and minimise the magnitude of the negative (leftward) displacement induced by relaxations? As in the figure, let us denote the pinch on the left ‘pinch 1’ and that on the right ‘pinch 2’. Pinch 1 produces maximum positive displacement (magnitude 4*L/*3, i.e. half of the optimal from Eq. (33)) when pinch 2 is completely open i.e. when the hydrodynamic resistance to its right is minimal. Pinch 2 then produces maximum positive displacement (of magnitude 8*L/*3, i.e. the optimal value in Eq. (33)) when pinch 1 is completely closed and all pinch-induced flow continues to be directed rightwards. Similarly, pinch 1 produces minimal negative displacement when pinch 2 is completely closed (zero average displacement, since no flow is then allowed to escape to the right of pinch 2), while pinch 2 then produces a negative displacement of magnitude 4*L/*3 to the left when pinch 1 is completely open. These optima can be achieved by the non-reciprocal sequence of motions illustrated in Fig. 5: close pinch 1, close pinch 2, open pinch 1, open pinch 2. The coordination between two pinches can therefore be used to generate the net displacement of 8*L/*3 equal to the theoretical upper bound from Eq. (33).

### J. Modelling of alternative flow generation mechanisms

We have described in detail our modelling of a network driven by the pinching of tubules. In this study we explore two other flow generation mechanisms: the contraction of tubular junctions and the contraction of ER sheets. Our model for pinching tubules is readily generalised to account for these mechanisms, as we describe below.

#### 1. Experimental estimates of junction volumes

From fluorescence microscopy images of ER networks, we measure the fluorescence intensity of junctions (i.e. the number of pixels in a junction multiplied by the mean intensity per pixel). We can then translate this intensity into an estimate of junction volume, assuming that intensity is directly proportional to volume. In order to calibrate the fluorescence intensity, we use our measurements for tubules. Specifically, we use the measured intensities of tubules and their known volumes, obtained by measurements of tubule lengths and the assumption that they are cylinders of radius 30 nm, in order to determine the proportionality constant to convert between pixel intensity and volume. Our new measurement of the intensities of the junctions then allows us to obtain estimates of their volumes; we obtain twelve values, ranging between 0.0020 μm^3^ and 0.0081 μm^3^, with a mean of 0.045 μm^3^ and a standard deviation of 0.0021 μm^3^.

#### 2. Mathematical modelling of contractions of tubular junctions

To include the contribution of tubular junctions into the model (Fig. 1c), we assume that in addition to the pinch sites along tubules, each tubular junction pinches independently of other tubules and other tubular junctions. Given a junction, we assume that it expels the same volume Δ*V* of fluid during a contraction for all its pinches (and takes in the same volume when it relaxes). Each pinch is assumed to create a sinusoidal flow source, so a pinch lasting for a time 2*T* produces a flow rate *S*(*t*) = Δ*V* sin(*πt/T*)*π/*2*T* at a time *t* measured from the beginning of the pinch, where the numerical factors in *S*(*t*) are chosen such that the volume expelled during a contraction is indeed 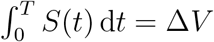. We accommodate the flows from the junctions mathematically by modifying our K1 equations, Eq. (17), to allow the normal nodes to also carry non-zero sources (as opposed to just the pinch nodes, as was the case before); the other equations in the model remain unchanged.

#### 3. Mathematical modelling of contractions of perinuclear sheets

In the tubules-pinching model, we included *M* exit nodes located towards the exterior of the network and through which flow could enter and exit the system in order to conserve mass. To account for the connection to a perinuclear sheet (Fig. 1f), we now assign randomly a number of these exit nodes, denoted *M*_2_ *< M*, to be ‘sheet nodes’ i.e. nodes which are directly connected to a perinuclear sheet, so that a number *M*_1_ = *M* − *M*_2_ *>* 0 of exit nodes remain. This is illustrated in Fig. 12, where we show the C1 network from Fig. 9 with both sheet nodes (blue asterisks) and exit nodes (red squares).

**FIG 12:**
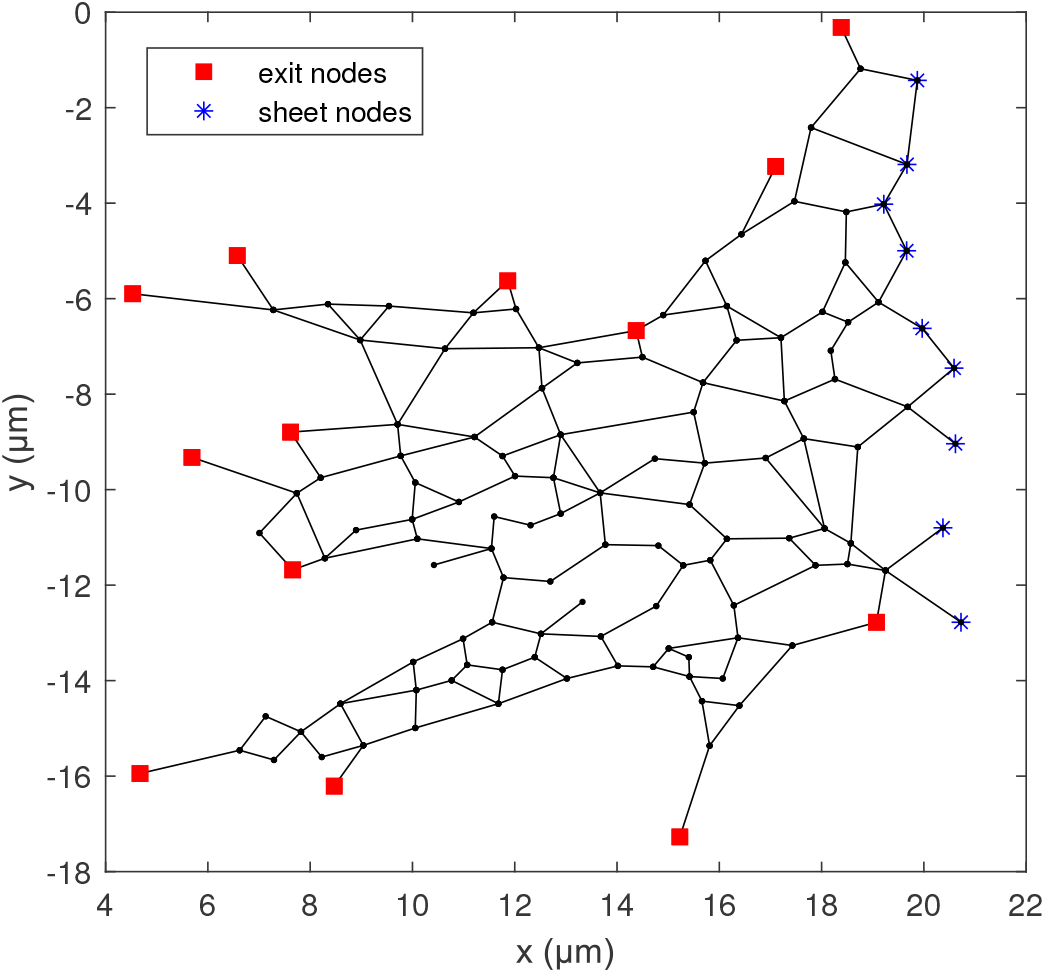
Illustration of the C1 network from Fig. 9 with *M*_1_ = 13 exit nodes (red squares) and *M*_2_ = 9 perinuclear sheet nodes (blue asterisks).

A sheet contraction+relaxation lasting a time 2*T* produces a total source *S*_sheet_(*t*) = *V*_sheet_*π* sin(*πt/T*)*/*2*T* at a time *t* from the beginning of the pinch, where again, the integral of *S*_sheet_ over a contraction gives a volume *V*_sheet_ of fluid expelled by the sheet.

Similarly to the mathematical model with tubular pinches only, our independent variables are the *E* tubule fluxes, the *M*_1_ sources at the exit nodes, and the *M*_2_ sources at the sheet nodes. As before, the K1 (Eq. 17) and K2 (Eq. 18) equations give us *E* + 1 equations. The requirement that the exit nodes are at the same (exit) pressure gives us 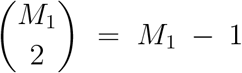 equations, namely the equality Δ*P*_*ij*_ = 0 for *i, j* exit nodes. We make the additional assumption that the sheet nodes are all at the same mechanical pressure (i.e. that they are connected to the same reservoir), which provides an additional *M*_2_ − 1 equations. Requiring that the sources at the sheet nodes sum to the prescribed flow rate *S*_sheet_ yields one additional equation, so we again have a system of *E* + *M* independent linear equations.

#### 4. Experimental estimates of volumes of peripheral sheets

To model the contraction of peripheral sheets (Fig. 1d), we need to estimate their contained volumes. Using the open-source image analysis software Fiji [38], we identify nine regions roughly occupied by peripheral sheets in a microscopy image of an ER network; this is illustrated in yellow in Fig. 13. We then measure their areas (in μm^2^) and convert them to volumes by multiplication with the diameter of a tubule, taken as 60 nm, assuming that the effective thickness of a sheet is equal to the tubular diameter. From our nine data values, we finally obtain a mean sheet volume of 0.12 μm^3^ and a standard deviation of 0.04 μm^3^.

**FIG 13:**
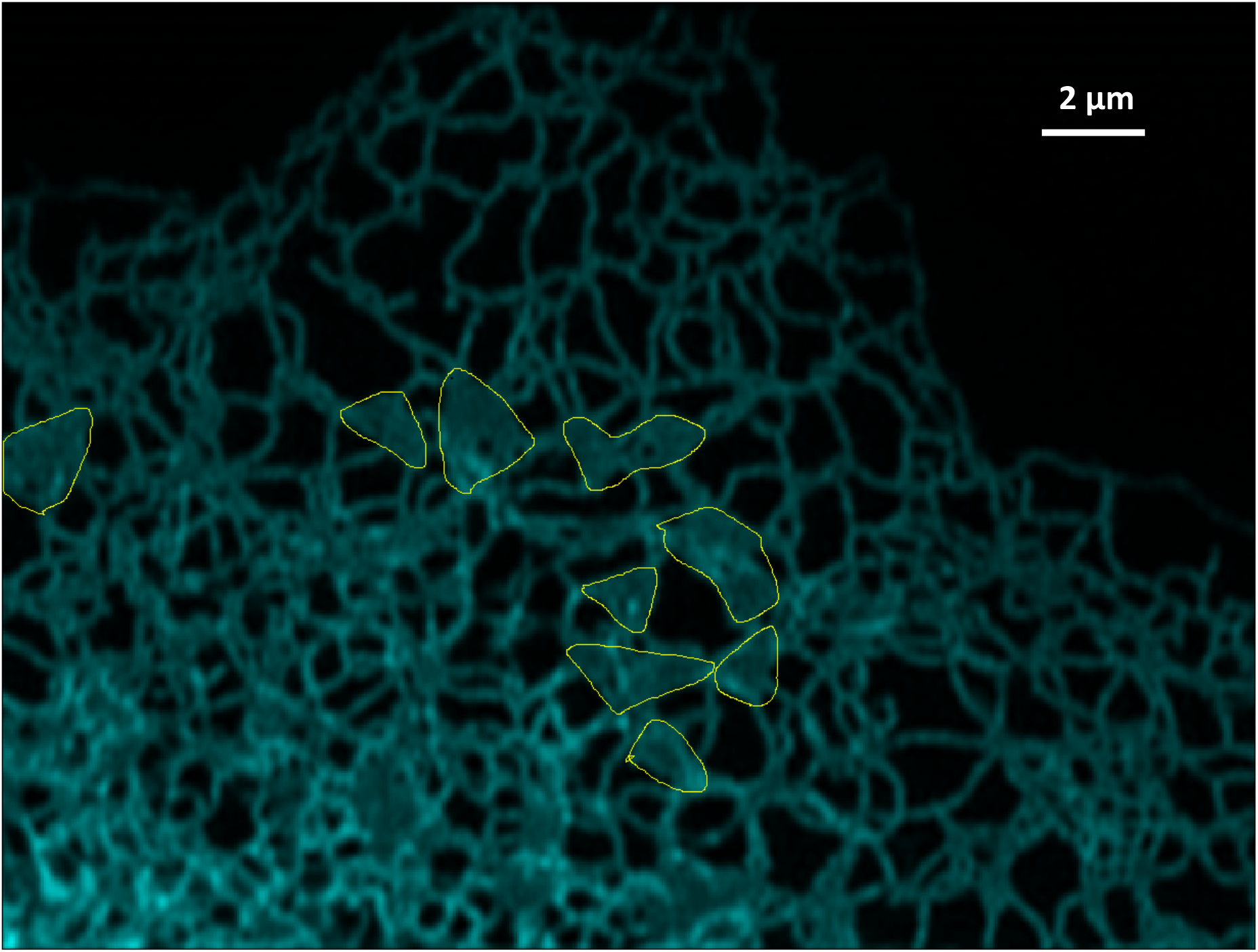
Estimation of areas of peripheral sheets, taken as the regions encircled in yellow.

### K. Energetic cost estimate for contracting peripheral sheets

The dominant energy expenditure in deforming a pinching tubule is in creating the large membrane curvatures at the narrowest sections of the pinch, and much less work is done against the small volumes of fluid displaced (§IV H). A contracting peripheral sheet, however, displaces a relatively large volume of fluid without attaining the extreme curvatures required in a pinching tubule. We therefore expect, intuitively, that the hydrodynamic contribution will dominate the energy expenditure budget. We now explicitly show this by estimating both elastic and hydrodynamic contributions.

To derive an order-of-magnitude estimate of the energetic cost, we consider an idealisation of a peripheral sheet consisting, in the relaxed state, of two parallel circular membranes *r < R* located at *z* = ±*D*, and in the fully contracted state, of two paraboloids at *z* = ±*Dr*^2^*/R*^2^ (see Fig. 14). The membranes deform as paraboloids between these two states, and we denote the ‘vertical’ distance between the two membranes as *d*(*r, t*). We take *R* = 0.8 μm, the value consistent with a sheet thickness 2*D* = 60 nm and the mean sheet volume (in the fully relaxed state) of 2*πR*^2^*D* = 0.12 μm^3^ (see estimation in §IV J 4).

**FIG 14:**
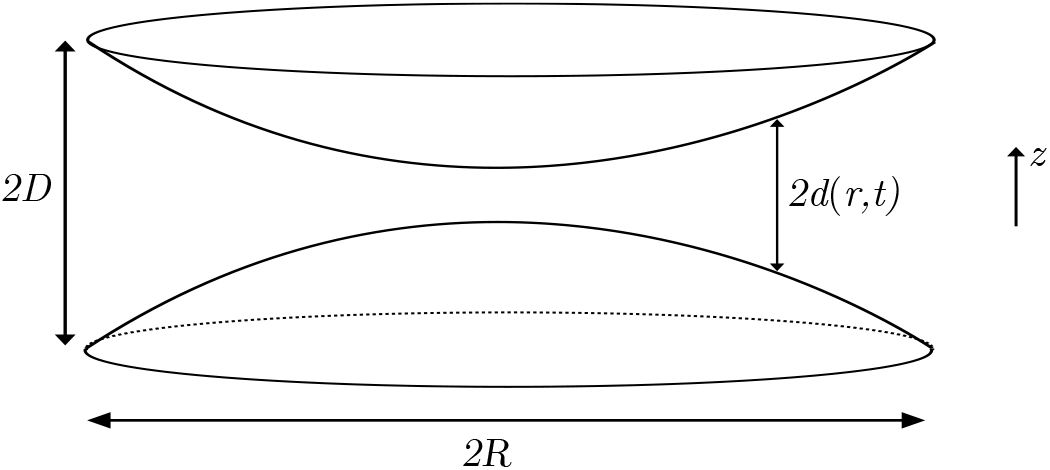
Mathematical idealisation of two contracting peripheral sheets as two paraboloids for the purpose of computing an order-of-magnitude estimate of the energy expended to contract a peripheral sheet.

To estimate the elastic contribution, we present an argument similar to the one carried out in §IV H. In the fully contracted state, the principal curvatures of one membrane scales as ∼ 2*D/R*^2^, so the Gaussian curvature *K* ∼ 4*D*^2^*/R*^4^ and the mean curvature *H* ∼ 2*D/R*^2^, yielding the Helfrich energy density *h* ∼ *k*[2*H*^2^ + *K*] ∼ 12*kD*^2^*/R*^4^, where *k* is the typical bending rigidity. The total bending energy of the fully contracted membrane is then *E* ∼ *πR*^2^*h* ∼ 2 × 10^−20^ J. In the relaxed state, the membranes are flat and have zero bending energy. Thus the total energy required to contract the two membranes can be estimated as

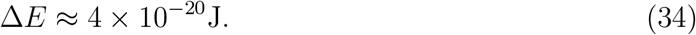

To estimate the hydrodynamic contribution, we first note that the work done instantaneously by the contracting sheet against the fluid, of dynamic viscosity *μ*, say, is the sum of dissipation rate in the sheet itself and the dissipation rate in the rest of the ER network outside the sheet due to the sheet-induced flows. Any work done against the fluid outside the ER network is neglected since flows decay over short length scales across the network.

We first consider the contribution from within the sheet. We may use lubrication theory to estimate the flows inside a contracting peripheral sheet since the sheet is fairly flat. The leading order lubrication flow between the contracting membranes (located at *z* = ±*d*(*r, t*)) is in the radial direction, and given by 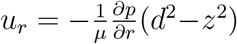, where the unknown radial pressure grathus yielding a radial velocity dient 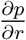 may be calculated from mass conservation 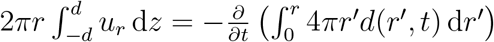, thus yielding a radial velocity

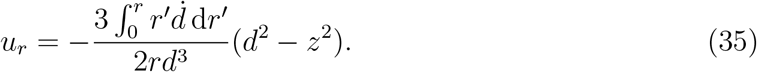

The dominant contribution to the rate-of-strain tensor is 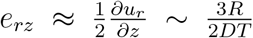, where *T* ∼ 0.5 s is the contraction duration (in the main text we considered a value of *T* of 2.5 or 5 times larger than the experimentally measured pinch duration; here we take 5). Therefore the total work done over a contraction scales as the total dissipation rate times the contraction duration *T*, and we scale

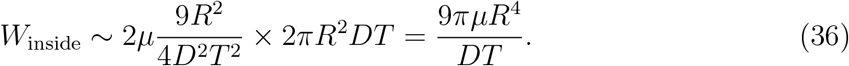

Taking the viscosity of the intraluminal fluid to be 10 times that of water, this is computed to be

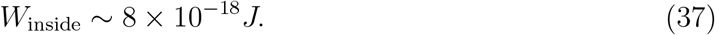

The dissipation rate in the network outside the sheet may be estimated by calculating the total dissipation rate in an idealised network consisting of ‘generations’ of tubules, each of length 1 μm, with the first generation consisting of three tubules connected to the peripheral sheet, and each tubule in the *i*^th^ branching out into three tubules of the (*i* + 1)^th^ generation ad infinitum. The dissipation rate due to a Poiseuille flow of flux *q* inside a tubule of length *l* and radius *R* may be calculated to be

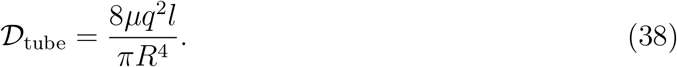

Denoting by *Q* the total volume flux created by the contracting sheet, we may calculate the dissipation rate in the network of tubules by summing the above expression across all the tubules as follows. The *i*^th^ generation of tubules is comprised of 3^*i*^ tubules each carrying a flux of *Q/*3^*i*^. Summing the total dissipation rates across all generations gives

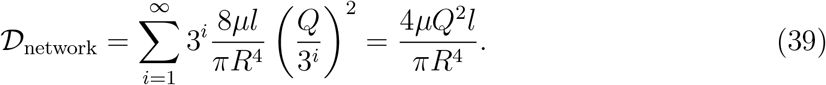

We may scale *Q* as *Q* ∼ *V*_s_*/*2*T*, recalling our assumption that each peripheral sheet contraction expels half of the volume *V*_*s*_ contained in the sheet (which is consistent with the paraboloidal membrane profile we have taken for the fully contracted sheet). The total dissipation over the contraction duration *T* in this network of tubules is therefore

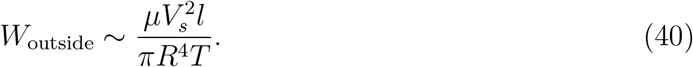

Scaling the sheet volume with its mean value *V*_*s*_ ∼ 0.12 μm^3^, we may compute this to be

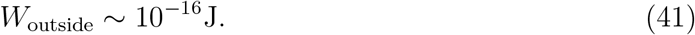

The total work done during a contraction is therefore *W* = *W*_inside_ + *W*_outside_ ∼ 10^−16^*J*. Given that the energy released by the hydrolysis of one ATP molecule is of the order of 10^−19^ J [39], we thus estimate that each contraction of a peripheral sheet would require on the order of 1000 molecules of ATP.

## Supporting information

Captions for Supplementary Videos

## ACKNOWLEDGMENTS

EL and PHH are supported by the European Research Council under the European Union’s Horizon 2020 research and innovation program (Grant No. 682754 to EL) and by the Cambridge Trust. EA is supported by the UK Dementia Research Institute [award number UK DRI-2004] which receives its funding from UK DRI Ltd, funded by the UK Medical Research Council, Alzheimer’s Society and Alzheimer’s Research UK

## Notes

### Competing Interest Statement

The authors have declared no competing interest.

### Summary of Updates

New subsection on energetic cost estimate of a peripheral sheet contraction; reordering and merging of some figures; various additional comments in main text

