## Supplementary material for "Fluid mechanics of luminal transport in actively contracting endoplasmic reticulum": Captions for Supplementary Videos

S1: Flows in an active C0 network pinching with the original pinch parameters. Edges are colour-coded with magnitude of instantaneous flow.

S2: Flows in an active honeycomb network pinching with the original pinch parameters. Edges are colour-coded with magnitude of instantaneous flow.

S3: Mixing in time in an a passive honeycomb network with no flow.

S4: Mixing in time in an active honeycomb network pinching with the original pinch parameters.

S5: Mixing in time in an active honeycomb network pinching with maximally long pinches at 10 times the original rates.
